# Preclinical lentiviral vector-mediated hematopoietic stem and progenitor cell gene therapy corrects Pompe disease-related muscle and neurological manifestations

**DOI:** 10.1101/2023.11.03.565442

**Authors:** John K. Yoon, Jeffrey W. Schindler, Mariana Loperfido, Cristina Baricordi, Mark P. DeAndrade, Mary E. Jacobs, Christopher Treleaven, Robert N. Plasschaert, Aimin Yan, Cecilia N. Barese, Yildirim Dogan, Vicky Ping Chen, Claudia Fiorini, Fritz Hull, Luigi Barbarossa, Zeenath Unnisa, Daniel Ivanov, Robert H. Kutner, Swaroopa Guda, Christine Oborski, Tim Maiwald, Véronique Michaud, Michael Rothe, Axel Schambach, Richard Pfeifer, Chris Mason, Luca Biasco, Niek P. van Til

## Abstract

Pompe disease, a rare genetic neuromuscular disorder, is caused by a deficiency of acid alpha-glucosidase (GAA), leading to the accumulation of glycogen in lysosomes and the progressive development of muscle weakness. The current standard treatment, enzyme replacement therapy (ERT), is not curative and demonstrates poor penetration into skeletal muscle and the central and peripheral nervous systems, susceptibility to immune responses against the recombinant enzyme, and the need for high doses and frequent infusions. To overcome these limitations, lentiviral vector-mediated hematopoietic stem and progenitor cell (HSPC) gene therapy has been proposed as a next-generation approach for treating Pompe disease. This study demonstrates the potential of lentiviral HSPC gene therapy to reverse the pathological effects of Pompe disease in a preclinical mouse model. It includes a comprehensive safety assessment via integration site analysis, along with single-cell RNA sequencing analysis of CNS samples to gain insights into the underlying mechanisms of phenotype correction.

**One Sentence Summary:** Preclinical hematopoietic stem cell gene therapy for the treatment of Pompe disease.

## Introduction

Pompe disease is a hereditary metabolic myopathy caused by a deficiency of the acid alpha-glucosidase (GAA) enzyme.(1, 2) It is characterized by the accumulation of glycogen in lysosomes, primarily in the heart, skeletal muscles, and central nervous system (CNS).(1–4) Infantile-onset Pompe disease (IOPD) patients have very low levels of GAA activity and experience severe muscle weakness. On the other hand, late-onset Pompe disease (LOPD) patients have higher residual GAA activity, leading to a slower disease progression, but often become dependent on wheelchairs and ventilators, with a reduced life expectancy.

The current standard of care (SOC) for Pompe disease is ERT which involves regular administration of recombinant human GAA (rhGAA).(5–7) While ERT can extend the lives of Pompe patients, it does not guarantee long-term symptom-free survival. The effectiveness of ERT is limited by several factors, such as poor penetration of rhGAA into affected muscle cells, low levels of mannose-6-phosphate (M6P) required for efficient cellular uptake, and abnormal M6P trafficking.

Lentiviral-mediated HSPC gene therapy has shown promise in treating other (neuro)metabolic diseases. Alternative developments to treat Pompe disease include adeno-associated virus (AAV) gene therapy and utilizing the hematopoietic system to produce rhGAA enzyme, which has been tested in preclinical models.(2) Unlike ERT, allogeneic HSPC transplantation has shown potential in promoting immune tolerance induction, which could enhance the efficacy of recombinant enzyme infusions.(8–10)

Additionally, liver-directed AAV gene therapy has also shown promising results in immune tolerance induction to rhGAA.(11) However, AAV serotypes targeting specific tissues, such as the liver, may not effectively deliver the transgene product to other affected areas, such as the CNS.

To enhance the efficacy of HSPC gene therapy, we employed a lentiviral vector containing a glycosylation-independent lysosomal targeting (GILT) tag sequence derived from the IGF2 peptide, which specifically binds with high affinity to the insulin-like growth factor 2 receptor (IGF2R) fused to a truncated catalytic domain of GAA. As we and others have previously demonstrated this fusion peptide improves enzyme secretion and enhances the reduction of glycogen, myofiber, and CNS vacuolation in critical tissues.(12–14) Additionally, the GILT-tagged vector sequence included an R37A mutein, which has been proven to significantly diminish signaling mediated by the insulin receptor.(14)

In this study, we have demonstrated consistent and elevated therapeutic enzyme activity, leading to complete or nearly complete reduction in the accumulation of disease-related substances in critical target tissues, including the CNS. Our analysis of integration sites revealed typical patterns associated with lentiviral vectors, without any indications of clonal expansions or biased selection of oncogenic sites. In addition, by performing single-cell RNA sequencing of microglia cells that were isolated using fluorescence-activated cell sorting (FACS), we observed that our therapeutic approach was capable of restoring these cells to a state similar to wild-type cells when compared to untreated glial cells or a non-GILT GAA vector. In summary, these findings underscore the potential of lentiviral HSPC gene therapy as a promising strategy for the treatment of Pompe disease, addressing the limitations of current therapeutic options.

## Results

### Hematopoietic reconstitution of genetically modified stem cells

In a previous study, a codon-optimized GILT-R37A-tagged GAA transgene (subsequently called *GILT*) was selected as a lead vector.(14) This vector contained the myeloproliferative sarcoma virus enhancer, negative control region deleted, dl587rev primer-binding site substituted (MND) promoter to drive *GILT* expression. In this study, to assess long-term expression and efficacy in *Gaa^−/−^* mice, we performed a lentiviral vector dose titration on *Gaa^−/−^* Lin-cells (multiplicity of infection (MOI): 0.75, 1.5 and 3), which were transplanted in both 9-12 weeks-old 7.5 Gy irradiated male and female *Gaa^−/−^* recipients. Supraphysiological plasma GAA activity levels were sustained up to the end of the study (week 31; Figure 1A). Peripheral blood (PB) leukocyte GAA activity was generally increased in the higher dose groups (MOI 1.5 and 3) compared to the lower MOI group (MOI 0.75) at week 4, 16 and 31 (Figure 1B and Suppl. figure S1A), as well as bone marrow at termination week 32 (Figure 1C). Using qPCR to determine vector copy number (VCN), PB leukocytes (male groups MOI 0.75 - 3, median per group 1.9 – 3.8 VCN and female groups 2.1 – 5 at week 4 and 1.9 – 3.3 and 1.8 – 3.4 at week 31 respectively), and bone marrow showed similar VCN patterns (male treatment groups median VCN 2.1 – 3.6 and females 1.9 – 3.6) (Figure 1D). The male MOI 3 group was often lower than the MOI 1.5 group (Figure 1D and Suppl. figure S1A) and the GFP group bone marrow VCN in males and females was 2.9 and 1.8 respectively. Using *in situ* hybridization assay to detect single nuclear lentiviral vector insertions, quantification showed a partial chimerism of vector positive bone marrow cells with a range of medians of 46% - 67% in males and 40 – 66% in females (Figure 1E). Quantification of the number of positive vector integrations per cell, showed that average nuclear single dots per bone marrow cell was slightly lower than quantified by qPCR (male treatment groups: median 1.3 – 2.1 and female treatment groups 1.1 – 2.2; Figure 1F). The average VCN in the transduced bone marrow cells <5, and the distribution of vector copies per cell similarly distributed between treatment groups and the GFP control group. The majority of vector positive cells contained a VCN of 1 (mean 33% - 49%), and cells with >5 VCN ranged from 9.2% - 13.9% in males and 10.6% – 15.6% in females. The MOI 3 GILT treatment group was used to determine VCN in peripheral tissues, and compared to the GFP group (MOI 1.5). In spleen and thymus, VCN was similar to PB leukocytes and bone marrow (VCN 2.8 – 3.3) in the GILT-R37A treatment group (Suppl. figure S1). In non-hematopoietic tissues lung, liver and kidney VCN were ∼3.5 – 17-fold lower. In reproductive organs testis and ovaries low VCNs were detected of 0.036 and 0.44 respectively (Suppl. figure S1).

**Figure 1.**
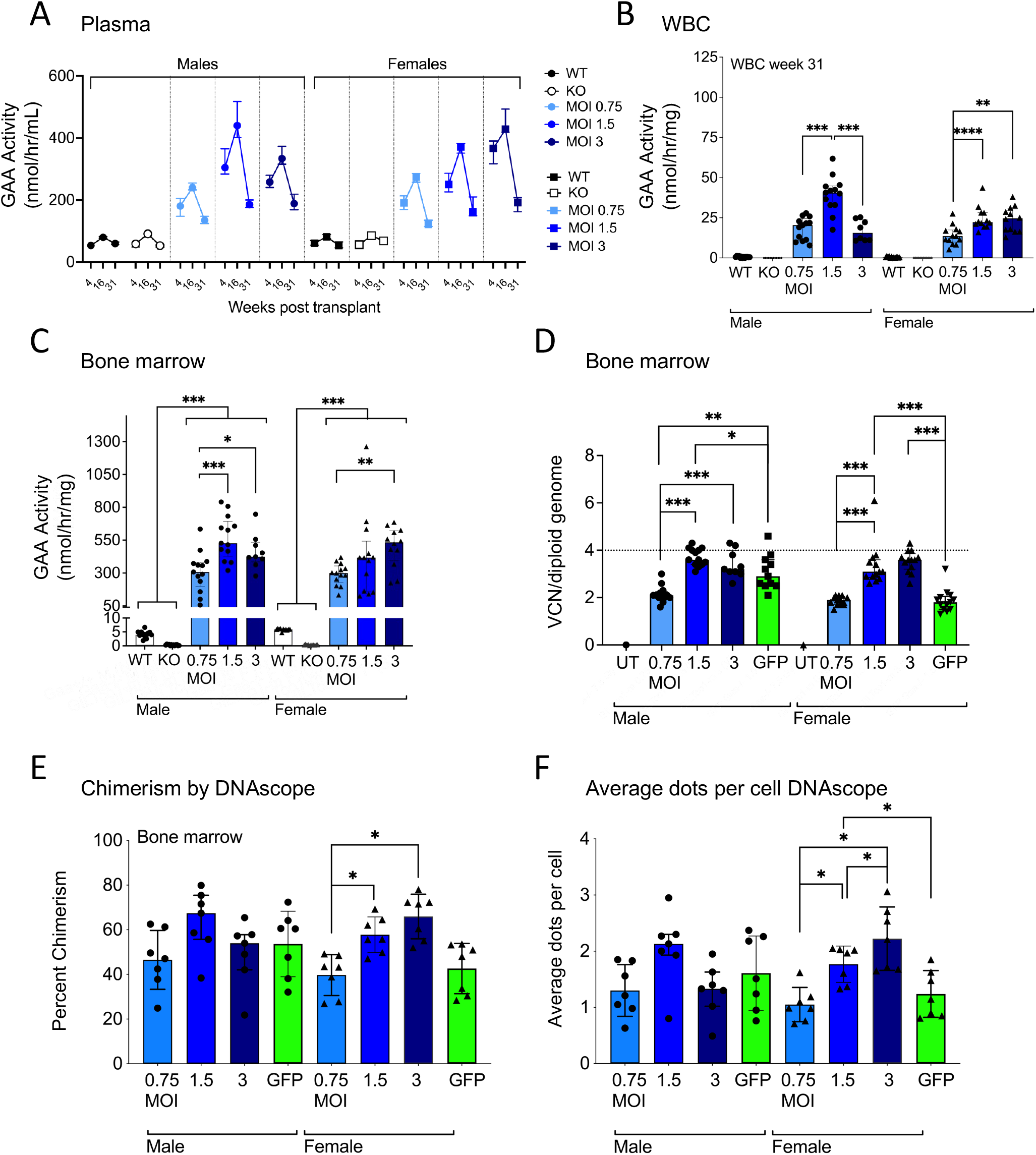
Hematopoietic reconstitution. (A) Enzyme activity in plasma at week 4, 16 and 31. (B) Enzyme activity in peripheral blood leukocytes at week 31. (C) Enzyme activity in bone marrow at 8 months after transplantation. (D) VCN/diploid genome in bone marrow determined by qPCR. (E) Chimerism based on nuclear detection of lentiviral vectors using DNAscope. (F) VCN quantified by nuclear vector-positive stained dots per cell using DNAscope. Bone marrow was collected 8 months after transplantation (N = 9 - 13). Statistical analysis; B-F: individual values, group medians, and interquartile ranges shown. Exact Wilcoxon Rank Sum p-values for group comparisons; *P <0.05, **P<0.01, ***P<0.001, ****P<0.0001.

### Complete correction of heart parameters

Heart GAA activity was restored to normal levels (median 3.8 in male and 4.5 in female *Gaa^+/+^*mice) and ranged from low MOI to high MOI 3.22 – 5.45 nmol/h/mg (male) and 3.1 – 7.7 nmol/h/mg (female) in the treatment groups (Figure 2A). Consequently, biochemical glycogen assessment showed complete elimination to undetectable levels (Figure 2B), which was confirmed by PAS quantification (Suppl. figure S2A, B). Cardiac remodeling and contractile function was affected in Pompe mice as previously described.(9) In our study, GILT-vector treatment resulted in normalization of heart weight (Figure 2C). Left ventricle (LV) mass and index was also determined by cardiac echography, and also showed reduction to normal values (Figure 2D, E). As for functional analysis, fractional shortening and ejection fraction were also significantly reduced after GILT vector treatment in all groups (Figure 2F, G), an indication of correction of hypertrophic cardiomyopathy. In addition, isovolumic contraction time (IVCT) was reduced in male recipient *Gaa^−/−^*mice (Figure 2H). Other cardiac parameters, such as LVVs, LVVd, LVPWs, LVPWd, IVSs, IVDd, LVIDs and LVIDd (abbreviations in Suppl. Table 1), which showed changes in size, mass geometry of the heart in Pompe mice, were improved and restored to normal ranges in GILT-treated mice (Suppl. figure S3A-H). Vacuolation scores also improved compared to male *Gaa*^−/−^ mice, that all had severity scores of 2, and treated groups all returned to normal in the MOI 3 group, and > 80% in the lower MOI groups (Suppl. figure S4). In the female groups, ∼70% had a severity score of 3, and > 80% of mice had no detectable tissue pathology after treatment. A low VCN was sufficient to drastically restore most cardiac parameters confirming the susceptibility to remodeling of the heart after GILT vector treatment.

**Figure 2.**
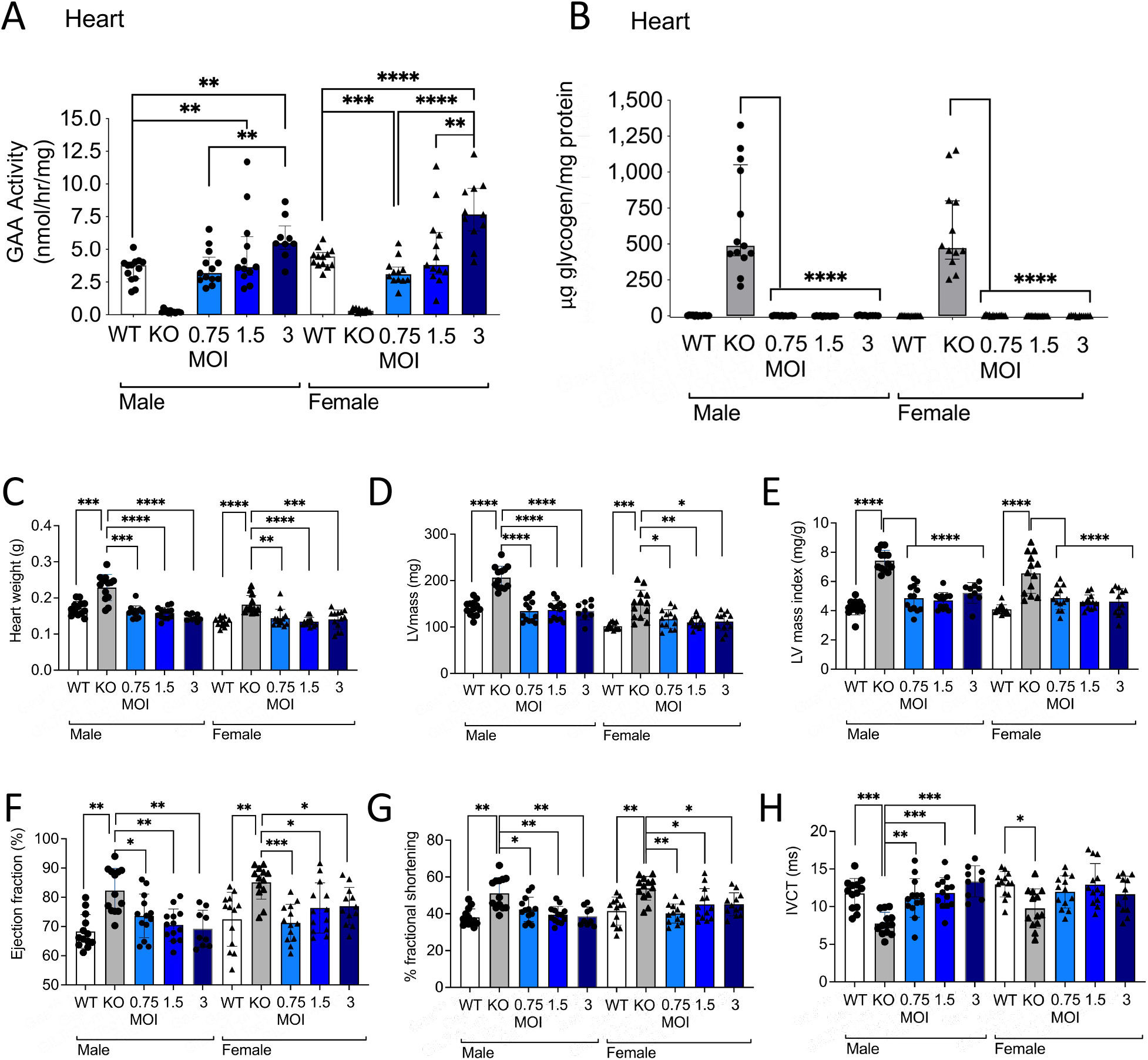
Cardiac function restoration. (A) GAA enzyme activity, and (B) glycogen in heart. Heart tissue was collected 8 months after transplantation (N = 9 - 13). (C) Heart weights (D) left ventricle (LV) mass (E) LV mass index, (F) Ejection fraction, (G) Fractional shortening (H), and isovolumic contraction time (IVCT). Analysis was performed 7 months after transplantation (N = 9 - 13). Statistical analysis; A-H: individual values, group medians, and interquartile ranges shown. Exact Wilcoxon Rank Sum p-values for group comparisons; *P <0.05, **P<0.01, ***P<0.001, ****P<0.0001.

### Correction of skeletal muscle pathology and improvement of locomotor function

The skeletal muscles diaphragm, as well as the hindleg muscles quadriceps, and gastrocnemius all consisted of higher GAA enzyme activities than the wildtype, in males 1.3 – 7.2-fold higher and females 1.3 - 7.9-fold higher (Figure 3A-D). In the tibialis only the highest MOI 3 resulted in a 2.3 fold increase in males and 1.9-fold increase in females. Consequently, glycogen was reduced > 98% in *Gaa^−/−^* treated male mice and >96% in *Gaa^−/−^* treated male mice in diaphragm (Figure 3E). In the hindleg muscles in *Gaa^−/−^*treated male mice, quadriceps, gastrocnemius (both >92%) and tibialis (>95%) glycogen was also significantly reduced (Figure 3F-H). In *Gaa^−/−^* treated female mice, quadriceps, gastrocnemius (both >85%) and tibialis (>93%).

**Figure 3.**
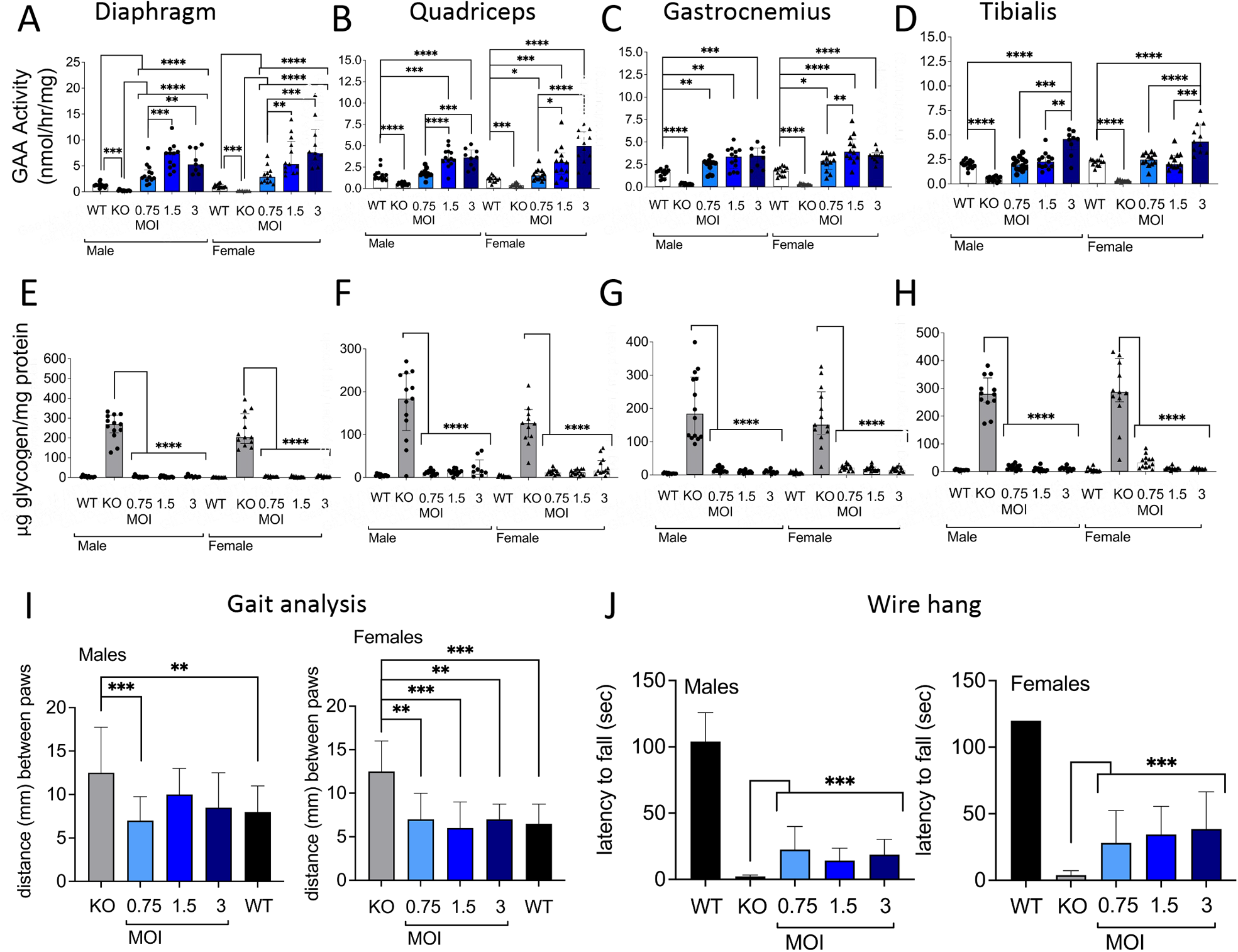
Skeletal muscle rescue. GAA activity in skeletal muscle diaphragm (A), quadriceps (B), gastrocnemius (C) and (D) tibialis muscle. Glycogen in muscle diaphragm (E), quadriceps (F), gastrocnemius (G) and (H) tibialis muscle at month 8 after transplantation (N = 9 - 13). Functional correction showing gait analysis (I) and wire hang at month 7 (J). Statistical analysis; A-H: individual values, group medians, and interquartile ranges shown. Exact Wilcoxon Rank Sum p-values for group comparisons; *P <0.05, **P<0.01, ***P<0.001, ****P<0.0001.

Quantification of PAS staining confirmed the glycogen content measured by the biochemical analysis of skeletal muscle lysates (Suppl. figure S2). Vacuolation scores in diaphragm males and females in the gene therapy groups were all lower than the *Gaa^−/−^* group, mostly with no detectable severity (Suppl. figure S4). In the hindleg muscles, the *Gaa^−/−^*always showed the most severe scores, and the gene therapy groups mitigation of skeletal muscle pathology with a trend in the low MOI group (MOI 0.75) containing more severe scoring compared to the high MOIs.

Locomotor function was also improved as assessed by gait analysis and wire hang, although the effects of gait were more pronounced in the female mice and the latency to fall was significantly increased, but back to normal in wildtype mice (Figure 3I, J).

### Correction of central nervous system pathology

In the CNS, cerebrum, cerebellum and spinal cord showed increases in GAA activity (median 5.4 – 28.2, 6.3 – 14.5 and 12.6 - 28.4 respectively in both males and females), in each case below the activities measured in wildtype mice (Figure 4A-C). However, glycogen was reduced more than 99% in all tissues (Figure 4D-F). Quantification of PAS staining confirmed the glycogen content measured by the biochemical analysis of CNS (Suppl. figure S2A and S2C). CNS tissue pathology was clearly detectable with vacuolation scores up to 5 in the *Gaa^−/−^* group (Suppl. figure 4). Remarkably, almost all gene therapy treated groups, including MOI 0.75, showed complete reduction and no detectable tissue pathology (Suppl. figure 4). VCN analysis in GFP-vector treated mice was similar to GILT-treated mice (Figure 4G). In a comparison study, we used GFP vectors to evaluate the strength of different promoter sequences *in vivo*. The MND promoter clearly showed the highest level of GFP mean fluorescence intensity in hematopoietic tissues including bone marrow, spleen, thymus and peripheral blood, compared to PGK and EFS regulatory sequences at similar VCN values (Suppl. figure S5A-E). In the CNS, the MND promoter was also robustly expressed in microglia-like cells, relative to moderate strength promoters EFS and PGK with similar VCN values (Figure 4H-J).

**Figure 4.**
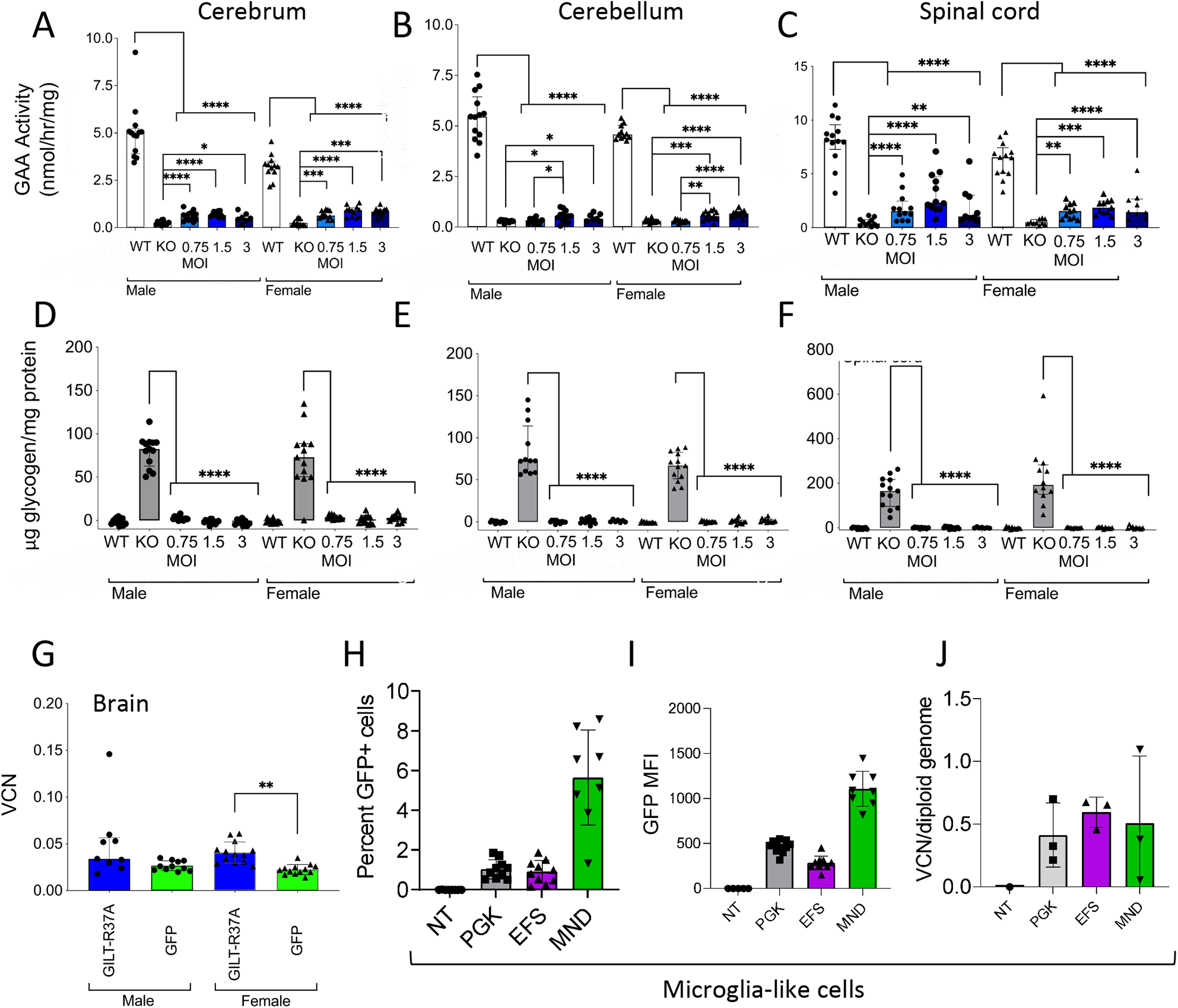
Restoration of function in the CNS. GAA enzyme activity in CNS, (A) cerebrum, (B) cerebellum, and (C) spinal cord. Reduction of glycogen in (D) cerebrum, (E) cerebellum, and (F) spinal cord. (G) VCN in brain. Tissues were analysed at 8 months after transplantation (N = 9 - 13). (H) Microglia cells isolated from mice transplanted with non-transduced (NT), and lentivector transduced Lin-cells containing EFS, PGK or MND promoter expressing GFP (N = 5-11). (I) MFI of GFP positive microglia cells. (J) VCN in brain. Tissues were analysed at 16 weeks after transplantation. Exact Wilcoxon Rank Sum p-values for group comparisons; *P <0.05, **P<0.01, ***P<0.001, ****P<0.0001.

### Brain single-cell transcriptional profiling

In order to evaluate the extent of phenotypic correction of microglia in the brain, spleen focus forming virus (SFFV) promoter driven lentiviral vectors SFFV.GILT or SFFV.GAAco vector were used to transduce Lin-cells, which were subsequently infused in Busulfex-treated mice. At 32-weeks after infusion, FACS-sorted CD45+CD11b+CXCR3+ microglia cells from the brain of the GILT vs GAAco treated groups (2 mice per group, 2 samples per mouse) were analyzed by single-cell(sc)RNA-seq (10x genomics). VCNs of Lin-cells, as well as in peripheral blood and bone marrow are shown in supplementary figure S6A-B. Immunohistochemistry for GAA protein revealed presence in the brain, but with distinct morphological patterns (Suppl. figure S6C). The GAA protein in the GAAco group was much more pronounced, compared to the GILT-group, which may be caused by the reduced ability of GAA protein secretion and cellular retainment of the GILT-protein as proposed by Dogan *et al*.(14) Immunophenotyping of single cell suspensions of glial cells showed an overall decrease in the percentage of microglia and increase in the proportion of astrocytes in untreated *Gaa^−/−^* and in GAAco-treated animals (Figure 5A and S6D). In contrast, the percentage of microglial cells and astrocytes was restored to WT levels in the GILT treatment group (Figure 5A). Endothelial cells and neurons were not affected (Suppl. figure S6E). Astrogliosis has been previously described in the Pompe mouse used in these studies.(15) In line with previous reports, we observed a higher percentage of astrocytes in nontreated *Gaa^−/−^* and GAAco treated animals, compared to WT animals. GILT treated mice had comparable percentages of astrocytes relative to WT animals. Microglial cells were sorted based on CD45, CX3CR1 and CD11b expression, and prepared for scRNA sequencing analysis (Suppl. figure 6D). We generated the combined single-cell transcriptional map (Figure 5B) where we identified 5 main clusters based on transcriptional similarities (see Material and Methods), which we annotated based on the differentially expressed genes reported in supplementary table S2. Notably, the microglia from GILT and GAAco mapped separately. The microglia from the GILT mice displayed an enrichment of clusters 1, 2 and 3 (generally corresponding to a more homeostatic defined signature) while the microglia from the GAAco treated group was highly concentrated in clusters 4 and 5 (reminiscent of disease-associated microglia signatures previously reported)(16–19) (Figure 5C). We observed higher expression of ribosomal genes in the microglia from GAAco while the GILT mice showed a statistically higher expression of genes normally associated with MHC complex expression, such as H2Eb1, H2-Ab1, Cd74 and H2-A. This phenotype was previously reported to be linked to border-associated macrophages which are generally yolk-sac-derived macrophages localized near the blood-brain barrier (Figure 5D).(20, 21) When looking for vector LTR presence in the mRNA of the single cells isolated from each experimental group, we could clearly observe a higher density of LTR+ cells in the microglia isolated from the GILT mice (Figure 5E), where ∼33% of the microglia were detected as vector positive (VCN 5.8, N = 2), versus 2.1% found in the GAAco group (VCN 2.4, N = 2). This near 16-fold higher level of LTR positive cells may indicate a preferential survival advantage for GILT-GAAco corrected microglia, which, as far as we know, has not been observed in other preclinical lysosomal storage disorders.

**Figure 5.**
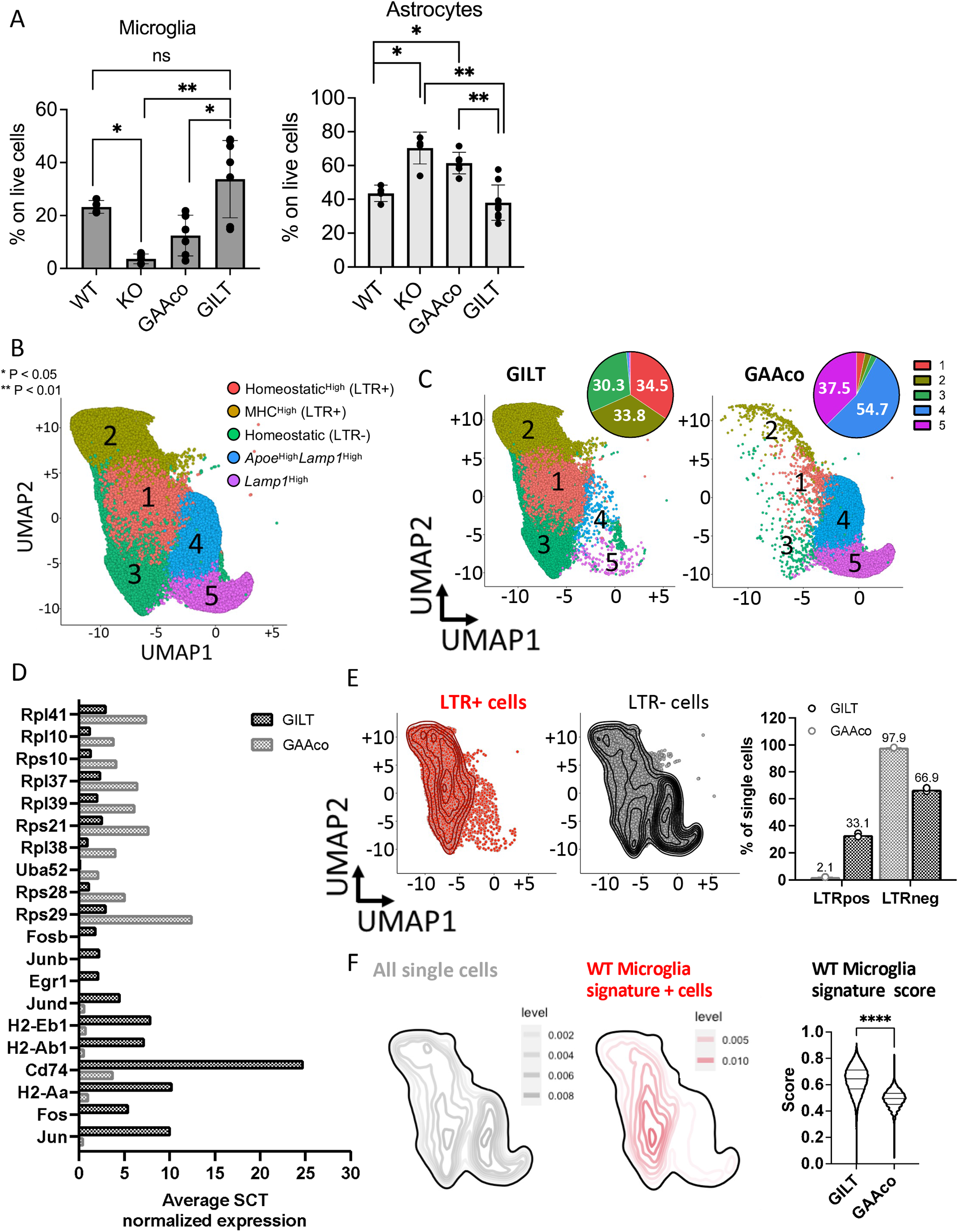
Single-cell RNA sequencing profile of microglial cells. (A) Percentage of microglial cells and astrocytes gated on live cells, measured via flow cytometry from enzymatically digested brain samples from mice 8 months after transplantation. (B) UMAP plot generated from a total of 53,421 single cells (N = 4 mice, 2 mice per group) clustered based on transcriptional similarities. Clusters are annotated on the top right based on their specific gene signatures listed in supplementary table S1. (C) UMAP plots showing single cell distribution in the GILT treated mice (left plot) or GAAco treated mice (right plot). Pie charts on the top right of each plot show the relative frequency of cells belonging to each cluster in the two groups. Percentages are reported for the most represented clusters in each group. (D) Bar plots displaying the average expression of ribosomal genes and homeostatic microglia-associated genes in microglial cells from GILT and GAAco treated mice. (E) Density plots showing the distribution of LTR+ (in red) and LTR- (in black) cells. The bar plot on the right show the percent of LTR positive (LTRpos) and LTR negative (LTRneg) in the GILT (dark gray bars) and GAAco (light gray bars) groups. (F) Density plots showing the distribution of all single cells in the map (in gray) vs the single cells expressing the WT microglia signature reported in supplementary figure S7D. WT microglia signature expression scores calculated in the GILT vs GAAco group are reported in the violin plot on the right. (Mann-Whitney-U p-values for group comparisons; ****P<0.0001).

Accordingly, analyzing the transcriptional profile of LTR+ vs LTR-cells (Suppl. figure S7) we could again identify gene signatures of activated microglia in the LTR-cells (more enriched in the GAAco group), while the LTR+ cells (more enriched in the GILT mice) showed a transcriptome more typical of homeostatic microglia. Based on these observations, we wanted to formally confirm that the cells isolated from the GILT mice carried a transcriptional profile more similar to normal microglia when compared to the GAAco group, as a result of a more efficient genetic correction driven by the GILT-vector. To this aim, we analyzed by scRNA-seq the microglia isolated from untreated wild type (WT) C57BL/6 mixed background control mice or Pompe mice (2 mice per group), generated the corresponding single-cell map (Suppl. figure 7B, C) and identified a list of genes specifically expressed by WT microglia (Suppl. figure 7D). When projecting this signature onto the GILT vs GAAco map we could clearly observe that the WT signature was highly enriched in the area of the map where the GILT cells were localized (Figure 5F). Accordingly, the WT signature score was statistically higher in the GILT vs the GAAco-treated mice suggesting that the GILT cluster contained a higher frequency of cells with a physiological microglia phenotype as compared to the GAAco group (Suppl. table 2).

### Genomic detection of integrated payloads and integration site analysis

We then applied our established protocol for insertion sites analysis (ISA)(22) combining our computational pipeline (23) to genomic samples from both irradiated and Busulfex-conditioned mice at 8-months and 16-weeks after infusion respectively, including GILT and GFP-vector treated groups. For the 8-months study, VCN per diploid genome of *in vivo* bone marrow samples from the GILT group ranged from 3.05 – 5.61. For GFP-vector group samples, the range was 1.90 – 5.33 copies.

In a separate *in vivo* experiment, male and female *Gaa^−/−^* mice were Busulfex-conditioned and subsequently transplanted with GILT lentiviral vector transduced Lin-bone marrow cells. Characterization of the transduction efficiency of the lentiviral vector lots used for these study are shown in supplementary figure S8A-C, as well as the GAA activity measured in WBCs and plasma at 4 and 12 weeks, in spleen and thymus at 8 (intermediate termination) and 16 weeks (Suppl. figure S8D). Hematopoietic reconstitution resulted in restoration of GAA activity in diaphragm heart and quadriceps (Suppl. figure S8E). Also, VCNs as in irradiated mice were similar in hematopoietic tissues and non-hematopoietic tissues (Suppl. figure S9A-B). Digital droplet PCR (ddPCR) was used to determine the VCN of *in vitro* samples (range 4.18 – 4.68 copies/genome) and of *in vivo* test samples from Group 2 (range 3.36 – 8.71 copies/genome). At takedown, GILT mice conditioned with either irradiation or Busulfex showed a polyclonal insertion site pattern as measured by the number of insertion sites retrieved and Shannon diversity. Results were similar between the GILT and GFP control group with no sign of aberrant clonal expansions in any of the mice analyzed (Figure 6A-D) (i.e., no integration in the GILT group contributed with more than 7.2 % or 6.32% to the total sequence pool for the irradiated mice or Busulfex conditioned mice respectively). For both the irradiation and Busulfex experiments, we did not observe any overlap among the Top 10 most abundant integration loci observed in each animal, suggesting that there was no obvious vector-driven selective growth associated to a specific gene locus (Suppl. figure S10). One animal had an integration inside *Kansl1l*, contributing with 20.84% to the overall sequence reads pool. Stocastic clonal selection is expected after transplantation. There was no direct link to hematopoietic malignancies regarding this gene. In the Busulfex experiment, the only reoccurring gene in the vicinity of a Top 10 integration was *Prkd1* for two animals. Of note, these are not the same chromosomal positions, but only the same gene. When looking at the insertion profile, we observed in both groups higher representation of integrations in a distance of 100-1000 kb relative to CpG islands, marking actively transcribed regions but not in the direct vicinity of CpG islands (1-10 kb) (Suppl. figure S11A). The vector generally disfavored GC-rich regions (100 bp – 10 kb, marking promoter regions of genes), longer gene widths or intergenic regions. These observations are in line with the classical insertional preference for gene bodies and underrepresentation of integrations in promoter regions described for lentiviral vectors (24). In addition, no selection for proto-oncogenic integration sites was observed *in vivo* in either of the groups (Figure 6D). Overall, these data suggest that regardless of the transgene and conditioning system used, our lentiviral vector displayed a polyclonal and typical insertional profile *in vitro* and *in vivo*.

**Figure 6.**
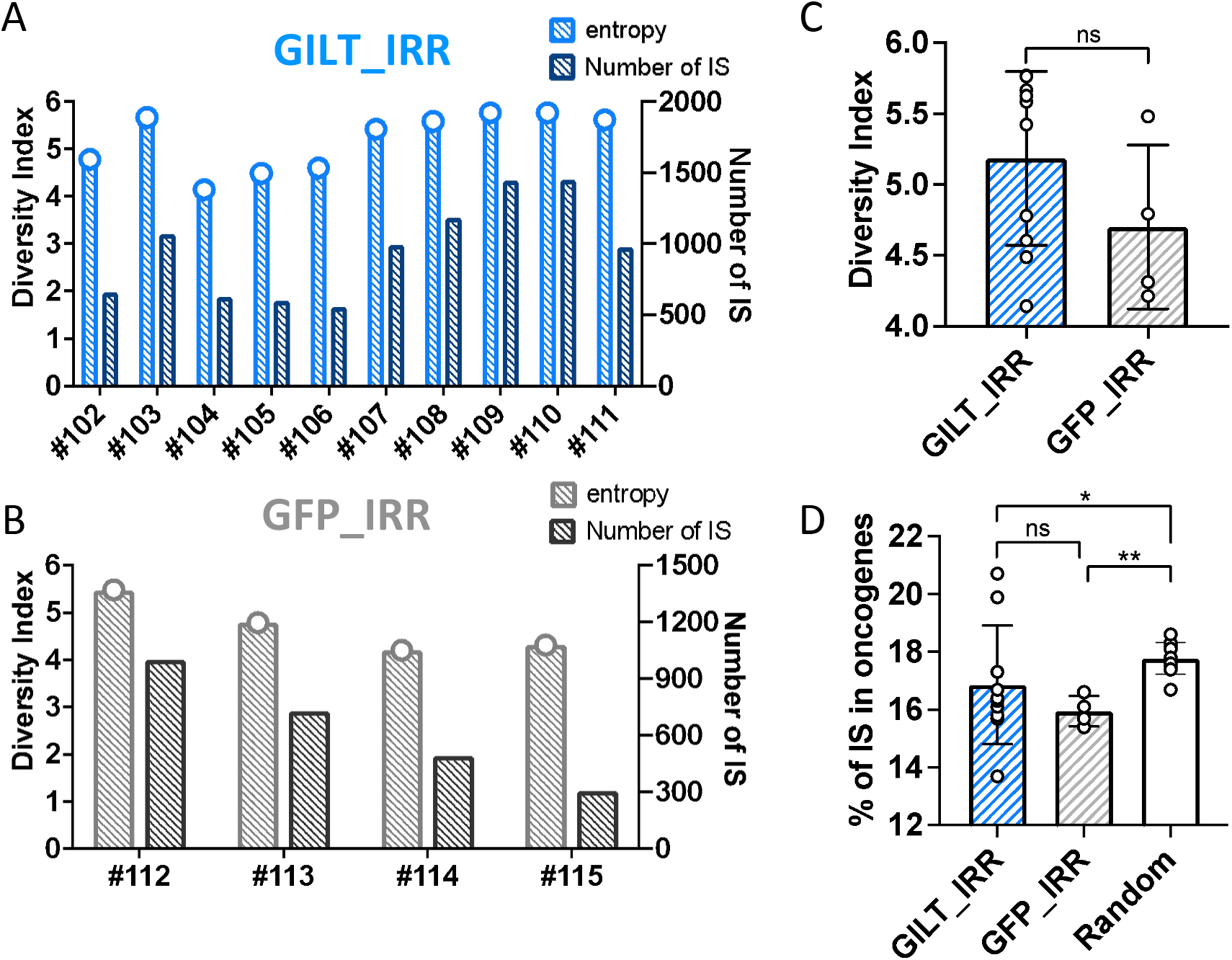
Integration site analysis. (A) Shannon diversity index and number of insertions collected at 8 months after transplantation in each mouse in the GILT (upper panel) or (B) GFP (lower panel) group from the study using irradiation conditioning. (C) Comparison of Shannon diversity indexes calculated on insertion sites collected from the GILT (blue bars) vs GFP (grey bars) group. (D) Percent of vector insertions detected in proto-oncogenes in the GILT group (blue bar), GFP group (gray bar) and a randomly generated insertion site dataset (white bar). Mann-Whitney-U p-values for group comparisons; *P <0.05, **P<0.01.

## Discussion

In this study, long-term efficacy and safety was observed in a mouse model of Pompe disease using a GILT containing lentiviral construct at low VCN. Heart parameters, which are relatively easy to normalize, compared to skeletal muscle, were shown to respond well to the therapeutic approach of using HSPC with GILT-technology in both male and female Pompe mice. Skeletal muscle involvement in Pompe disease is critical to improve disease pathology, but is more difficult to achieve with SOC. In our lentiviral approach, skeletal muscle parameters responded well after long-term gene therapy exposure, similar to what has also been observed in liver-directed AAV gene therapy.(25)

CNS involvement and pathology and contribution to phenotype has become more apparent in recent years, hence Pompe disease is generally considered a neuromuscular disorder. There are numerous reports that the CNS and peripheral nervous system (PNS) is associated with white matter abnormalities, cognitive impairment, and respiratory deficits. Subsequently the neurological component plays a critical role in disease progression, which is also not addressed by SOC.(3, 4, 26–30) Both the CNS and phrenic nerve dysfunction contributes to the respiratory insufficiency and failure, the most common cause of death in untreated IOPD and LOPD patients.(31) Moreover, neuropathology has also been investigated in Pompe mice, and double *Gaa^−/−^* transgenic mice expressing hGAA in skeletal muscle showed that skeletal muscle is not the sole contributor to ventilatory dysfunction.(32, 33) Furthermore, other investigators specifically investigated correction of neuropathology and respiratory dysfunction in Pompe mice.(25, 34, 35) An advantage of the followed approach using HSPC transplantation is that biodistribution of genetically modified cells is broad throughout the body and includes effective delivery to the CNS.(36) Furthermore, the MND promoter showed robust expression in microglia-like cells in the brain compared to two commonly used house-keeping gene promoters for HSC gene therapy, the EFS and PGK promoter. However, we only compared expression differences between the promoter variants with GFP as a model transgene. We did not investigate whether similar therapeutic effects could be obtained in the *Gaa^−/−^* mice using EFS- or PGK-driven constructs. We did not observe a major increase in VCN in the brain comparing GFP with GILT vector. Interestingly, isolation of microglia-like cells showed that GILT-treated mice had a major increase in the contribution of genetically modified cells. Of note, both astrocytes and microglia-like cell distribution appeared affected in Pompe mice, and the GAAco construct was unable to rescue CNS pathology, as well as microglia activation, which was normalized in GILT-vector treated mice.

Lentiviral HSC gene therapy has shown polyclonal integration profiles in metabolic diseases.(24, 37–39) In preclinical models for Pompe disease, lentiviral vectors with the strong SFFV promoter showed no significantly increased risk of in integration frequency near oncogenes compared to eukaryotic promoters.(40) A gammaretroviral vector with adenosine deaminase (*ADA*) cDNA under control of MND enhancer/promoter sequences has been used in clinical trials for ADA severe combined immunodeficiency (ADA-SCID). This demonstrated long-term findings with enduring efficacy, without developing malignancies, but also highlight risks of genotoxicity with gammaretroviral vectors, indicated by numerous common integration sites near proto-oncogenes and by increased abundance of clones with integrations near *MECOM* and *LMO2*.(41, 42) These clones showed stable behavior over multiple years and never expanded to the point of dominance or dysplasia.(42) In addition, the FDA-approved the product elivaldogene autotemcel (eli-cel or Skysona) for cerebral adrenoleukodystrophy (C-ALD), which uses a replication-incompetent lentiviral vector with *ABCD1* cDNA under the control of an internal MND promoter to transduce CD34+ HSPCs.(43–45) However, three patients treated with eli-cel in the ALD-102 (ClinicalTrial.gov ID: NCT01896102) and ALD-104 (ClinicalTrial.gov ID: NCT03852498) studies, were diagnosed with myelodysplastic syndrome (MDS) at 14 months, 2 years and 7.5 years after after Skysona administration (46, 47), and the hematologic malignancies appear to be a result of the Skysona lentiviral vector, Lenti-D, integration in proto-oncogenes (https://www.fda.gov/vaccines-blood-biologics/skysona). In our study, integration site analysis (ISA) showed typical lentiviral insertion patterns in transplanted mice conditioned with sublethal irradiation as well as Busulfex conditioning. Furthermore, genotoxic events were absent during the course of the study in HPSC transplanted mice containing GFP or GILT vectors, but this may also point out limitations to the sensitivity of testing vector genotoxicity in mice, and complementary assays such as IVIM/SAGA may aid to inform potential risk profile during lentiviral vector selection for clinical programs.(48)

Altogether, the *ex vivo* HSC gene therapy approach using the GILT-tag with R37A substitution demonstrated long-term efficacy in both CNS and skeletal muscle with a favorable safety profile in preclinical studies, which warrants further investigation.

## Materials and Methods

### Plasmid construction and lentiviral vector production

Lentiviral vectors containing the myeloproliferative sarcoma virus enhancer, negative control region deleted, dl587rev primer-binding site substituted (MND) promoter to express human codon optimized *GAA* (*GAA*co), and *GILT-R37A*-tagged *GAA* coding sequence were previously described.(14) In other experiments the spleen focus forming virus (SFFV) promoter was used for therapeutic transgene expression. Furthermore, lentiviral vectors expressing green fluorescent protein (GFP) were constructed with the following promoters: MND, phosphoglycerate kinase (PGK), and elongation factor 1 alpha short (EFS).

All third-generation self-inactivating LV vectors described above were produced by transient transfection of 293T cells using packaging and transfer plasmids at Vectorbuilder. Titers were determined on human osteosarcoma (HOS) cells (ATCC; Cat# CRL-1543) by qPCR and were subsequently titrated on lineage negative bone marrow cells to determine the MOI for subsequent studies.

### Mice

The *Gaa^tm1Rabn^*/J mice (*Gaa^−/−^* mice, Pompe mice) were used for the study.(49) Mice were obtained from an on-site breeding colony (kindly provided by Dr. Nina Raben, NIH, MD). Control B6129SF1/J mice (*Gaa*^+/+^ mice, WT mice) were obtained from Jackson Laboratories (Stock No. 101043). All mice were maintained in clean rooms and fed with irradiated certified commercial chow and sterile acidified water *ad libitum*. Assessment of animal health status, body weight, and examinations during the in-life term of the study were conducted by veterinarian personnel and documented. All protocols were approved by the Institutional Animal Use and Care Committee at the Canadian Council on Animal Care or the Charles River Accelerator and Development Lab (CRADL), Cambridge, MA.

### Assessment of dosing formulations and infusion in conditioned *Gaa^−/−^* mice

After *in vitro* assessment of vector preparations, *GILT*-tagged *GAA* containing lentiviral vectors were used for *in vivo* testing in male and female *Gaa^−/−^* mice mice. Bone marrow cells were harvested from femurs and tibias of 6-12-week-old male or female *Gaa^−/−^* donor mice, and Lin-enriched for HPSCs using RoboSep^TM^ (StemCell Technologies). After enrichment, cells were overnight transduced with each respective lentiviral vector, at a density of 10^6^ cells/mL in serum free StemMACS^TM^ medium containing 100 ng/mL recombinant murine stem cell factor (SCF); 50 ng/mL recombinant human FMS-like tyrosine kinase 3 ligand (Flt-3) and 10 ng/mL recombinant human thrombopoietin (TPO; StemCell Technologies). After sublethal conditioning of 9-12 weeks old *Gaa^−/−^* mice with 7.5 Gy gamma-irradiation (1.6 Gy/min) or by 4 x 25 mg/kg Busulfex^®^ (Otsuka Pharmaceutical) injections days −4 to −1 before cell dosing, the mice were injected intravenously with enriched donor Lin-HSPCs transduced with lentiviral vectors. Female *Gaa^−/−^* and *Gaa^+/+^* mice GFP vector and non-treated control groups were included.

Female or male *Gaa^−/−^* recipients that were irradiated were injected intravenously with 0.5×10^6^ cells per mouse from the opposite sex and monitored for 32 weeks with interim blood collections. Remaining transduced Lin-cells were cultured for VCN analysis. For Busulfex conditioned mice, same-sex Lin-cells were transplanted, and mice were followed up to 16 weeks after transplantation. At interim time points after cell infusion, leukocyte and plasma GAA activity, blood glucose, and complete blood counts using a Hemavet (Drew Scientific) analyzer were measured.

### Echocardiography Assessment

Complete echocardiographic examination was performed in all animals at month 7 after treatment (∼Week 24-28) at the Lady Davis Institute for medical research (Jewish General Hospital Sir Mortimer B. Davis, 3755 chemin de la Côte Sainte-Catherine, Montréal, Québec, H3T 1E2), using a FUJIFILM VisualSonics echocardiography system (Vevo 3100, FUJIFILM VisualSonics, Inc. Canada). The echocardiography results were evaluated by the technical specialist at the Lady Davis Institute.

Animals were transferred to the treatment facility in a transport vehicle with controlled temperature. Sterile transport bins were modified to allow up to 8 animals/bin with food pellets and gel packs. Clinical signs were monitored immediately before and after transportation. At the test site, the animals were placed in dorsal recumbency on a heated surface and allowed to reach steady-state resting heart rate (500-550 bpm) prior to echocardiography examination.

Echocardiography was performed from a left parasternal approach using standards from American Society of Echocardiography for measures. Analyses were done using M-mode and pulse wave Doppler signal with a short axis view and an apical two chamber view. Echocardiography was performed as described previously with some modifications.(50, 51) Briefly, the mice were anesthetized with 3.5% isoflurane mixed with O_2_ at 1 L/min. The depth of anesthesia was confirmed by the rear foot squeezing. The mice were secured lightly on supine position on a warming pad adjusted to 38°C, and their anterior chest was shaved. The echocardiography was performed under isoflurane anesthesia using a VEVO 3100 ultrasound machine and a MX550S transducer with a center frequency of 40 MHz (FUJIFILM VisualSonics Inc., Toronto, ON, Canada). The body temperature and heart rate (HR) were maintained respectively to 36-37°C and 500-550 beats/minute during the study. Body temperature was measured using a rectal probe and maintained by adjusting the intensity of an infrared lamp that is positioned over the mouse. HR could be influenced by changes in body temperature, with lower HR with temperature below 36°C and higher HR with temperature above 37°C. Furthermore, even if body temperature was within the require range, mice could present bradycardia. This was corrected by decreasing the % of isoflurane up to 2%. However, if HR increased too much, the % of isoflurane was re-increased gradually up to 3.5%.

Left ventricle (LV) systolic function was assessed as follows. Two-dimensional guided M-mode images were obtained from a short axis view at the papillary muscle level to determine the LV structure and systolic function. The LV internal diameter in diastole (LVIDd) and the interventricular septum and LV posterior wall thickness in diastole (IVSd and LVPWd) were measured. LV mass, LV mass index and fractional shortening (FS) were calculated as follows.

LV mass (mg) = [(LVIDd + IVSd + LVPWd)^3^ - LVIDd^3^] x 1.055 x 0.8, where 1.055 is the density of the rat myocardium (in mg/mm^3^) (17) and 0.8 a correcting factor to compensate for the overestimation of LV mass. LV mass index (mg/g) = LV mass/body weight. FS (%) = [(LVIDd + LVIDs)/LVIDd] x 100. LV mass index (LVMi) was determined as follows: LVMi (mg/g) = LV mass x 0.8/body weight.

### Locomotor function tests

The gait analysis was performed to evaluate the locomotor function in mice. For that the entire underside of all toes and the center of the feet of the animal were fully covered in paint. After that mouse was placed on a piece of white paper at the start of the white plastic corridor and allowed to walk all the way into the black box (safety place). At least two steps consistently spaced with clear, non-smudged footprints were measured from each foot for fore-to-hind paw distance. The male animals were tested before the female animals for any testing day. The test was performed as per standard operating procedure (SOP) including habituation runs on 2 occasions for the groups on month 6 and 7.

The wire hang test was performed to assess the muscular strength of mice. For that the animal was placed on the cage top, which was then inverted and suspended above the home cage. The latency until the animals fall was recorded. The average performance for each session was presented as the average of the three trials. The test was performed as per SOP on 3 occasions for all animals in month 7.

### Blood glucose monitoring

Blood glucose was measured as previously described (14) at different time points during the in-life study and prior to scheduled termination with glucometer Accu-chek AVIVA. Animals were fasted overnight prior to collections.

### Scheduled termination procedure

Scheduled termination was either weeks 16 or 32 post-transplant. Animals were fasted 8-12 hours prior to euthanasia and anesthetized with isoflurane. After cardiac puncture blood collection, the animals were perfused with phosphate-buffered saline (PBS) pH 7.4 until internal organs were pale in appearance. The heart mass was weighed, and the dissected tissues were snap frozen and stored at −80°C for GAA activity and glycogen measurement or processed for histopathology. Therapeutic endpoints included biochemical GAA activity, tissue glycogen content, VCN analysis, histological evaluation, blood glucose, and immunohistochemistry in brain sections.

### Measurement of GAA enzyme activity and glycogen by biochemical analysis

For GAA enzyme activity and glycogen measurement, tissue samples of heart, diaphragm, gastrocnemius, quadriceps femoris, tibialis anterior, cerebellum, cerebrum were collected at necropsy and processed chilled to homogenization in sterile dH_2_O, centrifugation, removal of clear supernatants, and then stored at −80°C until the selected assays were performed. Similarly, terminal plasma and cell pellets from peripheral blood, spleen and bone marrow samples were collected and kept at −80°C for GAA enzyme activity.

The GAA enzyme activity was measured similar as described.(14) Study samples were assayed in a 96 well plate using fluorescent synthetic substrate 4-methylumbelliferyl-alpha-D-glucosidase (4-MU) at 6 mM concentration, in presence of 9 µM acarbose and 90 minutes incubation. A 0.5 M carbonate buffer pH 10.7 was used to stop the reaction and assay plates were read at 365 nm excitation and 450 nm emission in SpectraMax^TM^ i3X (Molecular Devices, CA). This method was qualified over the range of 0.1 to 81 nmol/mL. Results were normalized for protein concentration in the sample using Bicinchoninic Acid (BCA) kit (Pierce, ThermoScientific).

The glycogen in tissues was estimated treating samples with and without *Aspergillus niger* amyloglucosidase that generates β-D-glucose which is oxidized to release gluconic acid and hydrogen peroxide.(14) In presence of horseradish peroxidase, hydrogen peroxide reacts with o-dianisidine hydrochloride to generate a colored product detectable at 540 nm in a spectrophotometer and is proportional to the glucose concentration in the sample. The qualified assay had a dynamic range of 5.6 to 160 µg/mL. Results were normalized for protein concentration in the sample using BCA kit (Pierce, ThermoScientific).

### VCN analysis

VCN quantification was completed either by quantitative PCR (qPCR) or digital droplet PCR (ddPCR). All qPCR assays consisted of oligonucleotide primers and probe mixes containing either a TaqMan 6-carboxyfluorescenin (FAM) or VIC^TM^ fluorescent probe designed to amplify the *HIV Psi* vector sequence or housekeeping genes glycosyltransferase like domain containing 1 (*Gtdc1*) or transferrin receptor protein 1 (*Tfrc*) as previously described.(14, 36) For qPCR VCN assessment, a single plasmid containing both sequences was used as a reference standard in a range of 50 to 5 × 10^7^ copies. Data was reported as VCN/diploid genome.

For ddPCR, specific primers targeting the *HIV Psi* element were used to detect the integrated lentiviral vector, along with specific primers targeting genomic reference sequences *Gtdc1*.

### Immunohistochemistry

Brain hemispheres of GAAco and GILT-vector treated mice were perfused fixed in 4% formaldehyde overnight at 4°C, then washed 3X and stored in PBS. The specimens were embedded, and arranged for coronal sectioning in a gelatin matrix using MultiBrain^®^ Technology (NeuroScience Associates, Knoxville, TN). After curing with a formaldehyde solution, the blocks were rapidly frozen by immersion in 2-methylbutane chilled with crushed dry ice and mounted on a freezing stage of an AO 860 microtome. The MultiBrain^®^ blocks were sectioned coronally with a setting on the microtome of 35µm. All sections were cut through the entire cortex of the brain and collected sequentially into a series of cups. All cups contained Antigen Preserve solution (50-parts PBS pH7.0, 50-parts ethylene glycol, 1-part polyvinyl pyrrolidone).

For immunohistochemistry, selected sections were stained free-floating. All incubation solutions from the primary antibody onward used Tris buffered saline (TBS) with Triton X100 as the vehicle; all rinses were with TBS. After a hydrogen peroxide treatment and rinses, each section was immunostained with primary antibody overnight at room temperature. Following rinses, the primary rabbit antibody against GAA (ab240102 Abcam) and secondary goat anti-rabbit biotinylated (Vector Laboratories, BA-1000) was applied. After further rinses Vector Lab’s ABC solution Catalog # PK-6100 (avidin-biotin-HRP complex for VECTASTAIN^®^ Elite ABC, Vector, Burlingame, CA) was applied. The sections were again rinsed, then treated with a chromagen: diaminobenzidine tetrahydrochloride (DAB), nickel (II) sulfate and hydrogen peroxide to create a visible reaction product. Following further rinses, the sections were mounted on gelatin coated glass slides, air dried, then dehydrated in alcohols, cleared in xylene and coverslipped. Brightfield stainings were scanned using a Huron Digital Pathology LE120 TissueScope. Slides were scanned using a 20x objective for a final image resolution of 0.4 μm/pixel.

### Periodic Acid-Schiff and H&E staining of mouse tissues

Following scheduled necropsy, tissues from *Gaa^−/−^* and *Gaa^+/+^* mice were divided for different endpoint analyses, and a portion of each collected, preserved, trimmed, and placed in separate cassettes. Periodic Acid-Schiff (PAS) and hematoxylin and eosin (H&E) stainings were performed as described.(14) Tissues were fixed in 10% NBF for up to 32 hours and post-fixed in 1% periodic acid (PA)/10% neutral buffered formalin (NBF) for 48 hours at 4°C. One hemisphere from each brain sample along with heart, diaphragm, tibialis anterior, gastrocnemius, and quadriceps femoris tissues were fixed in 10% NBF for up to 32 hours and further post-fixed in 1% PA/10% NBF for 48 hours at 4°C and processed into FFPE blocks. Blocks of cerebral cortex, cerebellum, hippocampus, and/or brainstem, thoracic and cervical spinal cord, heart, quadriceps femoris, diaphragm, gastrocnemius, and tibialis anterior were sectioned at 4 microns, mounted onto glass slides, and stained with PAS and H&E, and evaluated for glycogen accumulation and vacuolation by light microscopy. Tissues stained via PAS/H protocols were scanned at 20X magnification using a Hamamatsu Nanozoomer whole-slide scanner. Scan files were imported into Visiopharm for quantitative image analysis. A region of interest (ROI) surrounding individual tissue sections was automatically applied to all scan files using a Visiopharm Analysis Protocol Package (APP). Automated ROIs were manually refined to optimize anatomic homology across animals. Automated image analysis APPs were used to detect the dark purple PAS+ signal contrasted against a pink PAS-background. The following equations were performed in Visiopharm to generate quantitative end-points: 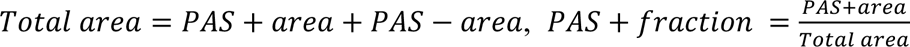, and 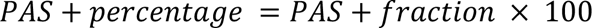. Quantitative immunofluorescent data were exported from Visiopharm as excel spreadsheets.

Vacuolation in CNS tissue was limited to specific cell types and sub-anatomic sites. Therefore, severity grading was assigned based on the estimated number of visible neurons with abundant cytoplasmic vacuoles within the most affected anatomic regions (e.g., brain stem, spinal cord gray matter, sciatic nerve). Scores were assigned as: N (vacuolation absent); 1 (< 10% of neurons heavily vacuolated); 2 (≥10% and up to 25% of neurons heavily vacuolated); 3 (≥25% and up to 50% of neurons heavily vacuolated; or less than 25% total with heavily vacuolation of specific brain stem nuclei, e.g., gracile nucleus), 4 (≥ 50% and up to 75% of neurons heavily vacuolated); 5 (≥ 75% of neurons heavily vacuolated). Vacuolation within myofibers in muscle (cardiac, skeletal; smooth muscle) was scored as: N (vacuolation absent); 1 (minimal vacuolation visible throughout section, requiring at least 20x microscope objective to confirm vacuoles were present); 2 (vacuolation clearly visible with a 10x objective, vacuoles are limited within individual myofibers and do not form large aggregates); 3 (vacuoles clearly visible at 10x and lower objectives with aggregates forming around nuclei and center portions of myofibers); 4 (large vacuolation aggregates are prominent in many myofibers); 5 (most myofibers contained large vacuolation aggregates).

### DNAscope for nuclear detection of lentiviral copy number

RNAscope^®^ Assay is an RNA *in situ* hybridization (ISH) approach that allows simultaneous signal amplification and background suppression to achieve single-molecule visualization while preserving tissue morphology.(52, 53) The approach was modified to visualize lentiviral vector DNA (DNAscope®). DNAscope® target retrieval conditions were optimized for detection of single lentiviral vector DNA using oligoprobes in bone marrow cells of treated mice. DNAscope® pretreatment conditions were as follows: Epitope Retrieval (LS ER2): 15 minutes at 88°C, Protease III: 5 minutes at 40°C, Customized denaturation DNAscope® Protocol DNA *in situ* hybridization for detection of DNA was performed on the Leica automation platform using the RNAscope® 2.5 LS Red Reagent Kit (Advanced Cell Diagnostics, Inc., Newark, CA) along with custom accessory reagents according to the manufacturer’s instructions. Briefly, samples on cytospin slides were pretreated with heat and protease prior to hybridization with the target oligo probe. Preamplifier, amplifier, and alkaline phosphatase-labeled oligos were then hybridized sequentially, followed by chromogenic precipitate development. Specific DNA staining signal was identified as red punctate dots. Samples were counterstained with Hematoxylin. Slides were scanned with the 3DHistech Panoramic SCAN II digital scanner to perform HALO^®^ image analysis to detect and quantify lentiviral vector positive cells throughout the entire bone marrow cytospin preparation and copy number measurement as binned dots per cell.

### Integration site analysis

Samples from both *in vivo* studies were used to perform integration site analysis. Bone marrow isolated from 16 transplanted male and female *Gaa^−/−^* mice (n = 14 with the GILT-vector and n = 2 with non-transduced cells) at 8 months after transplantation from 7.5 Gy irradiated mice. Additionally, 8 *in vitro* samples (cultured lineage negative cells), seven days after transduction with GILT-vector (two samples each of MOI 0.75, 1.5, and 3) or MND.GFP (two samples of MOI 1.5) lentiviral vectors. Mice were selected for ISA based on the highest VCN/diploid genome (dg) measured at week 12 and week 16 in the peripheral blood DNA (determined by qPCR).

The second study included the bone marrow samples obtained from male and female *Gaa^−/−^* mice from the Busulfex study (n = 5/sex) with transduction was done at an MOI of 5. The mice were culled at 6-weeks after Lin-bone marrow cell transplantation. Three *Gaa^−/−^* mice administered non-transduced Lin-bone marrow cells (male n = 1; female n =2) were also included in this analysis as controls.

INSPIIRED sample processing was performed as described in Sherman *et al*., 2017,(22) and adapted and described in Ha *et al*, 2021.(54) DNA samples in nuclease-free water were subject to shearing (Covaris 220) for 60 seconds at a peak power of 50 Watts, 5% duty factor, 200 cycles/burst, 4°C water temperature. Samples were AMPure purified (0.7-fold bead to sample ratio) and used for end preparation of fragmented DNA using the NEBNext Ultra End Repair/dA-Tailing Module. Previously generated linkers (linker blunt + sample-specific linker) were introduced with the NEBNext Ultra Ligation Module. Samples were again AMPure purified (0.7-fold bead to sample ratio) and used in exponential PCR1 with linker-specific primers (300nM), and LTR specific primer 1 (300nM), blocking oligo (1 μM), dNTPs (200 nM), Clontech Advantage PCR buffer (1x) and mix (1x) (TaKaRa), linker-ligated DNA (0.6 v/v ratio), water (0.172 v/v ratio) and a thermocycler program of 1 min at 95°, and then 20 cycles of 30 sec at 95°C, 30 sec at 80°C and 90 sec at 70°C for exponential amplification and finally 4 minutes at 72°C. For the second nested PCR, the primers, blocking oligos, dNTPs, Clontech Advantage PCR mix, and buffer had the same concentration as in PCR1, and the exponential amplification step in PCR2 was performed with 15 cycles. For all reactions, specific index primers and sample-specific linker primers were used. Afterwards, PCR2 reactions were mixed in equal volumes to generate the final library 210303_INSPIIRED_RUN40. The library was first column purified prior to two AMPure purifications with a 0.7-fold, and 0.6-fold ratio of beads to sample volume. Libraries were transferred to the research core unit genomics (RCUG) of Hannover Medical School for quality control via Bioanalyzer and analysis by Illumina sequencing on flow cells with 15 million clusters.

Bioinformatic analysis was generally performed as described by Berry and colleagues.(55) The analysis files necessary to run the INSPIIRED pipeline were downloaded from GitHub (https://github.com/BushmanLab/INSPIIRED). Individual sequence files were aligned and annotated to the mouse genome (mm9). The plasmid vector sequences served as a reference for LTR regions and vector trimming. The processing and alignment statistics were exported before uploading the results to a local database using inSiteUploader.R. The data frame with all integration site data was exported and used for customized post-processing steps in Microsoft Excel 2016, GraphPad Prism (Version 5) and R (3.6).

### Isolation of microglia/MLCs and single cell RNA-Seq

After transcardial perfusion, the brains of four female mice, two from SFFV.GILT and two from the SFFV.GAAco experimental groups were processed by enzymatic digestion (Neural Tissue Dissociation Kit (P), Miltenyi Biotec). The resulting single cell suspensions were stained with viability dye and antibodies for microglia cells, neurons, astrocytes and endothelial cells (LIVE/DEAD Fixable Aqua Dead Cell Stain Kit, ThermoFisher; CD45.1 clone A20 in BV421 (BD); CD45.2 clone 104 in BV421 (BD); CD11b clone M1/70 in APC-780 (ThermoFisher); CX3CR1 clone SA011F11 in BV605 (BioLegend); PECy7 Thy1.1 (also known as CD90.1) clone OX-7 in PE-Cy7 (BD); Thy1.2 (also known as CD90.2) clone 53-2.1 in PE-Cy7 (BD); ACSA-2 clone IH3-18A3 in PE (Miltenyi Biotec); CD31 clone 390 in PerCP/Cy5.5 (BioLegend). Microglia cells, defined as CD45+ CD11b+ Cx3cr1+, were FACSsorted using a MA900 Multi-Application Cell Sorter (Sony Biotechnology). For the generation of single cell transcriptomes, a target cell number of 2.5×10^3^ or 5×10^3^ cells from each sorted population were run using the Chromium Controller (10x Genomics) using Chromium Next GEM Single Cell 3’ Reagent Kits (10x Genomics). The libraries generated were then run on the NextSeq 550 (WT vs Pompe microglia datasets) or the NovaSeq 6000 (GILT vs GAAco datasets) Sequencing System (Illumina) using either the using NextSeq 500/550 High Output v2.5 (150 cycles) Kit or the NovaSeq 6000 S2 Reagent Kit v1.5 (100 cycles) Kit (Illumina). The Illumina raw BCL sequencing files were processed through the CellRanger software (10x Genomics) for generating FASTQ files and count matrixes (https://support.10xgenomics.com/single-cell-gene-expression/software/overview/welcome). which were then used as input for the SEURAT V4.0 (https://satijalab.org/seurat/) R tool for single cell genomics analyses. Briefly, single cell barcodes were filtered for the ones containing mitochondrial gene content lower than 15%. Expression data then were normalized, scaled, and searched for variable features using the SCTransform function of SEURAT V4.0 followed by UMAP dimensionality reduction and clustering using the *FindClusters* function with resolution set at 0.2. The maps shown in figure 5B,D and supplementary figure S7B,C were generated using the *ggplot* RPackage based on the UMAP coordinates from the SEURAT package. The differentially expressed genes shown in figure 5C and supplementary figure 7A,D were identified through the *FindMarkers* function of SEURAT and plotted either using Prism 9 (GraphPad Software, LLC.) or through the *pheatmap* R package (https://cran.r-project.org/web/packages/pheatmap/pheatmap.pdf). The signature scores of figure 5F have been calculated through the *UCell* R package (https://github.com/carmonalab/UCell) using as signature the WT-specific genes shown in supplementary figure 7D.

### Statistical analysis

Statistical analysis was performed using either Prism (GraphPad) or R. For scRNA-Seq data we used GSEAPreranked analysis to identify the gene sets that are enriched in LTR-negative and LTR-positive samples. To perform this analysis, we firstly derived a score using log2FoldChange and p-value from differential gene expression analysis (score=-s*log10(p-value) with s=-1 if the log2FoldChange of gene and p-value from differential gene expression analysis (score=-s*log10(p-value) with s=-1 if the log2FoldChange of gene. For all other data, a Student t-test (when only two groups were present), one-way ANOVA with Tukey’s post hoc analysis (when multiple groups were present), or linear regression was used. Significance was defined as p < 0.05.

Experimental groups were sized to allow for statistical analysis; not all the animals were included in the analysis, and select outliers were excluded. Mice were assigned randomly to experimental groups based on weights. All other statistical analysis was analyzed by VERISTAT, Inc or internally by applying median and interquartile range using Kruskal-Wallis and exact Wilcoxon rank-sum tests for groups comparison and SAS/STAT^®^ software v9.4. A result of < 0.05 was indicative of significant difference in the groups. For correlation analysis, the Pearson R coefficient and p-value were used.

Quantitative immunofluorescent data were collected in Visiopharm and statistical analysis was performed using SAS software (v9.4). Continuous variables were analyzed via Levene’s test for equality of variance. Where Levene’s test was not significant, one-way ANOVA was used to detect differences among three more group means. Tukey’s test was used to explore pairwise group comparisons when one-way ANOVA was significant. Where Levene’s test was significant, the Kruskal-Wallis test was used to detect differences between three or more group means. The Dwass-Steel-Critchlow-Fligner method was used to explore pairwise group comparisons when Kruskal-Wallis test was significant. Significance was set to p < 0.05 for all statistical tests. In the figures *P<0.05, **P<0.01, ***P<0.001 and ****P<0.0001.

## Acknowledgements

We thank all the members of AVROBIO, Inc. for their continued support of our work. We would like to thank Charles River Laboratories for study execution. Additionally, KCAS Bioanalytical & Biomarker Services for their contributions to sample analysis, NeuroScience Associates (NSA) for the immunohistochemistry of brain, and VERISTAT, Inc for biostatistical analysis. Finally, we would like to thank Nina Raben (National Institutes of Health, MD) for supplying the *Gaa* knockout mice.

## Author contributions

JY, JWS, CB, contributed to the experimental design and execution, biochemical and molecular assays development and qualification, data analysis and interpretation, and wrote the manuscript; ML, MPD, MEJ, CT, RNP, AY, YD, CB, VC, CF, FH, LB, ZU, DI, RHK, SG, VM, MR, AS conducted experiments, and performed data analysis, MR and AS contributed to experimental design and analysis; TM performed statistical analysis, CO, RP, CH contributed to assay development and study logistics; TM performed statistical analysis; CM contributed to the design and the reporting of the work presented; LB, NvT contributed to the design, data analysis, interpretation and writing of the manuscript.

## Data availability statement

Data supporting the studies presented in this manuscript can be found in the main text or the supplemental information. Additional information, as appropriate, may be made available by request directed to the corresponding author and, as appropriate, following execution of a suitable confidentiality agreement with AVROBIO, Inc.

