## Supplementary Materials for "Preclinical lentiviral vector-mediated hematopoietic stem and progenitor cell gene therapy corrects Pompe disease-related muscle and neurological manifestations"

### Supplementary figures

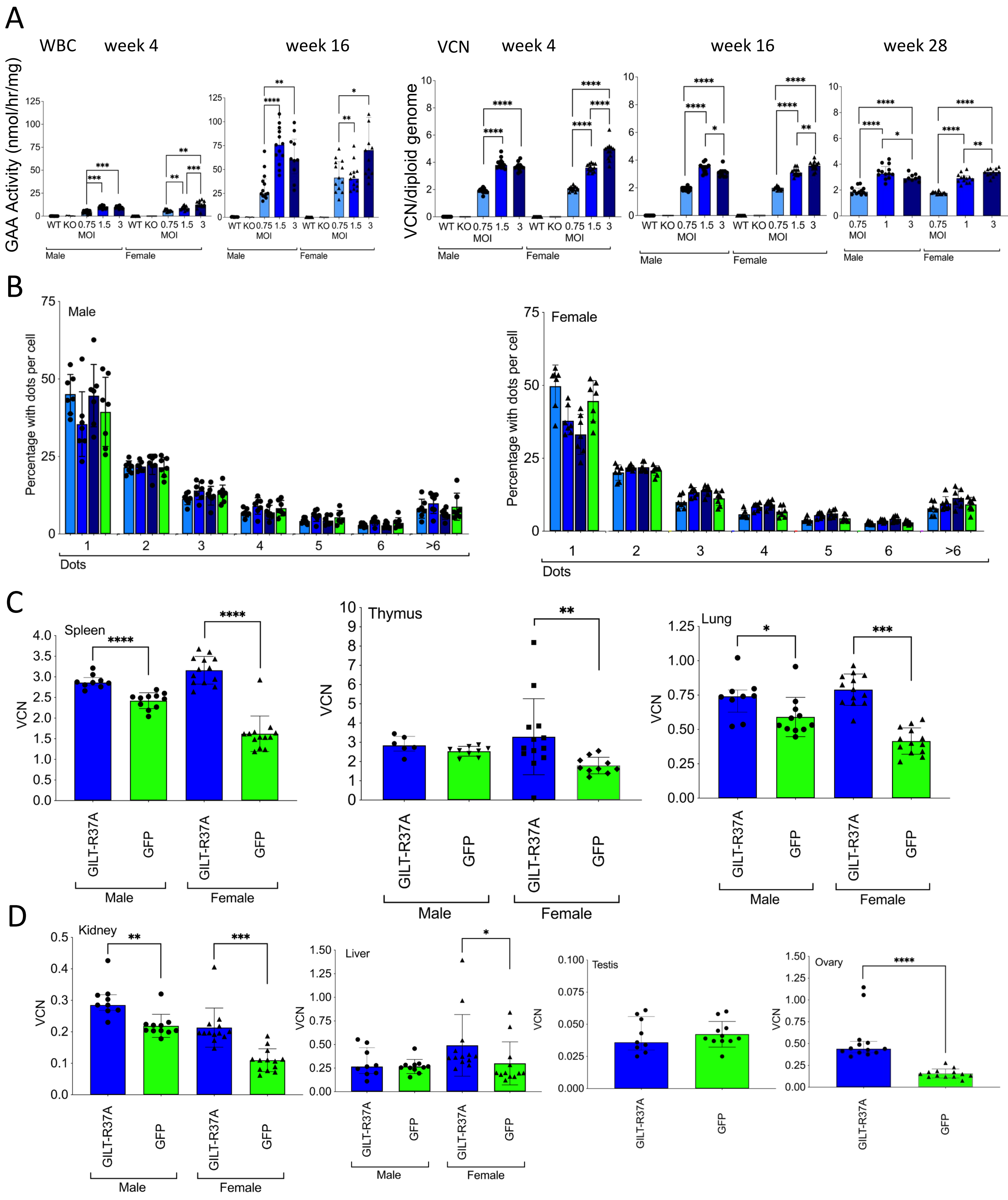


**Supplementary figure S1.** Reconstitution in the hematopoietic compartment and peripheral tissues. (A) GAA enzyme activity week 4 and 16, and VCN in peripheral blood white blood cells. (B) Quantification of dots per cell by DNAscope technology. Distribution in males and females. Mean +/- SD (C) VCN in hematopoietic tissues spleen and thymus and other peripheral tissues lung, kidney, liver, testis and ovary. Statistical analysis (N = 6-13); A, C, D: individual values, group medians, and interquartile ranges shown. Exact Wilcoxon Rank Sum p-values for group comparisons; *P <0.05, **P<0.01, ***P<0.001, ****P<0.0001. VCN = vector copy number, WT = wildtype, KO = knockout.

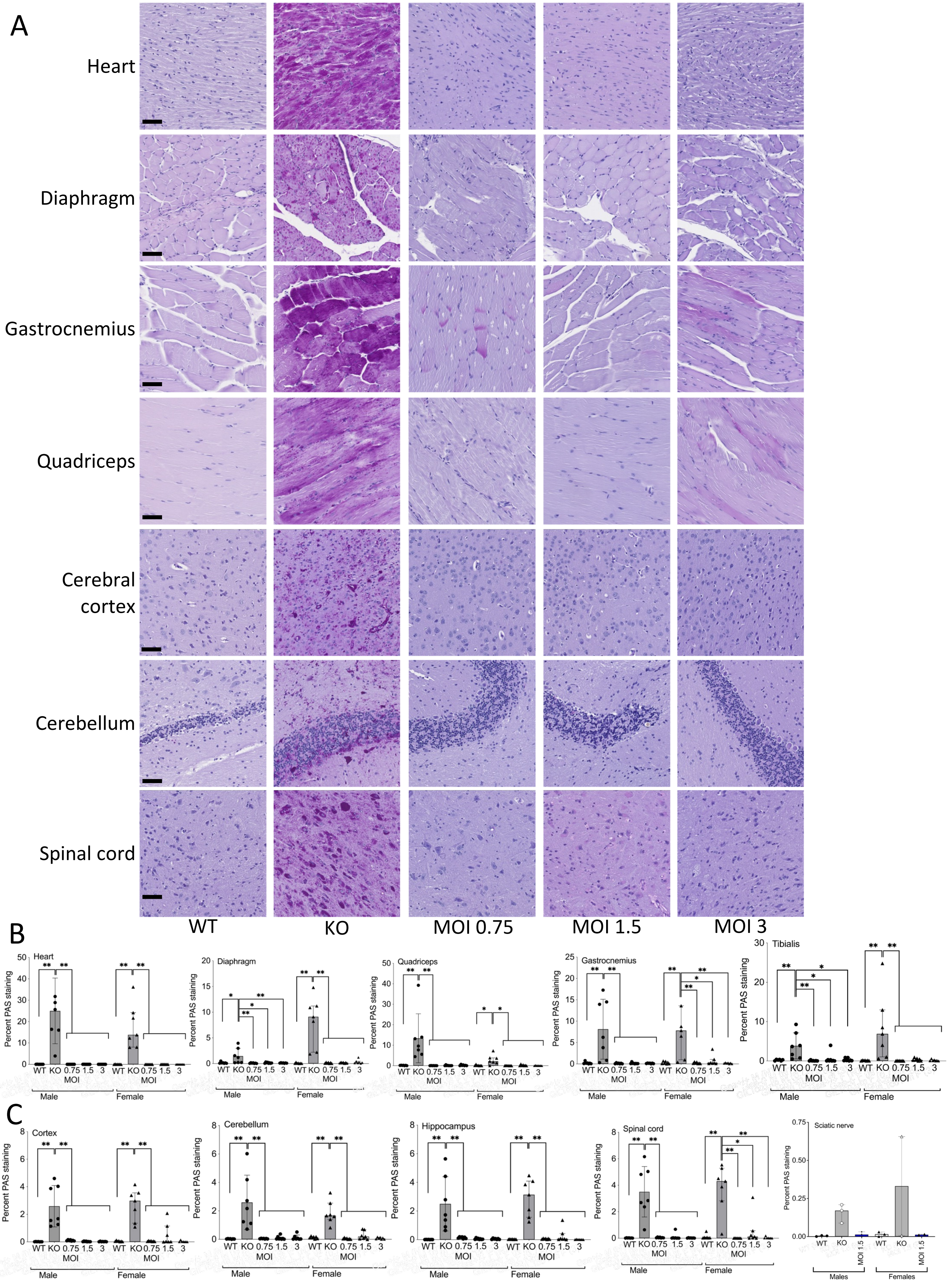


**Supplementary figure S2.** Glycogen clearance in GILT-treated *Gaa*$^{-/-}$ mice. (A) PAS staining of muscle tissues and CNS of female treated *Gaa*$^{-/-}$ mice (scale bar 50 μm). (B) Quantification of glycogen in muscles (C) Quantification of glycogen in CNS. Statistical analysis (N = 7-8); B, C: individual values, group medians, and interquartile ranges shown. Exact Wilcoxon Rank Sum p-values for group comparisons; *P <0.05, **P<0.01, ***P<0.001, ****P<0.0001. CNS = central nervous system, WT = wildtype, KO = knockout.

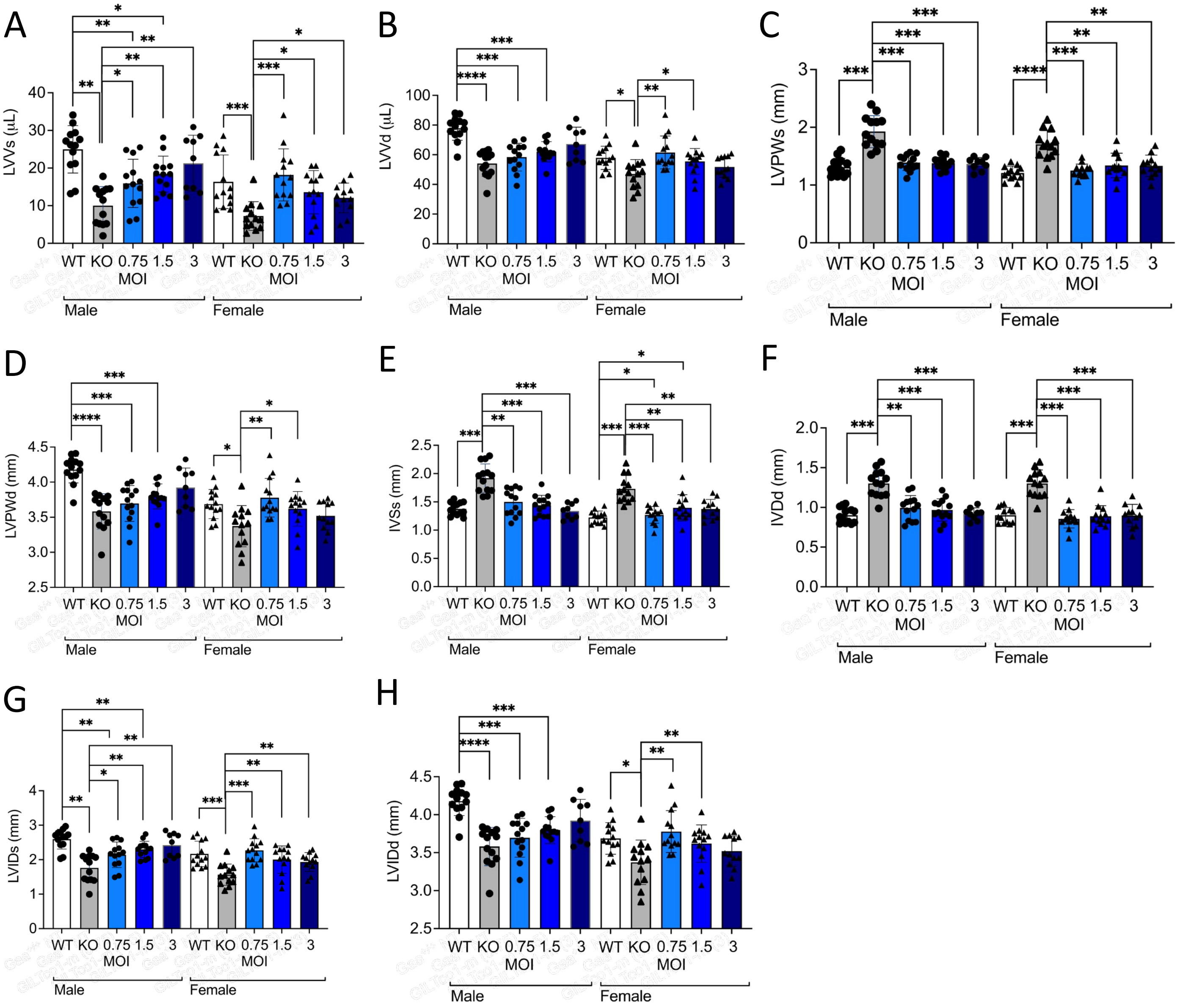


**Supplementary figure S3. Echocardiography.** Functional rescue of cardiac parameters measured by cardiac echography. (A) Left ventricular volume at systole (LVVs) (B) Left ventricular volume at diastole (LVVd) (C) Left ventricular posterior wall at systole (LVPWs) (D) Left ventricular posterior wall at systole (LVPWs) (E) interventricular septum thickness at systole (IVSs) (F) interventricular septum thickness at diastole (IVSd) (G) left ventricular inner dimension at systole (LVIDs) (H) left ventricular inner dimension at diastole (LVIDd) Statistical analysis (N = 9-13); A-J: individual values, group medians, and interquartile ranges shown. Exact Wilcoxon Rank Sum p-values for group comparisons; *P <0.05, **P<0.01, ***P<0.001, ****P<0.0001. WT = wildtype, KO = knockout.

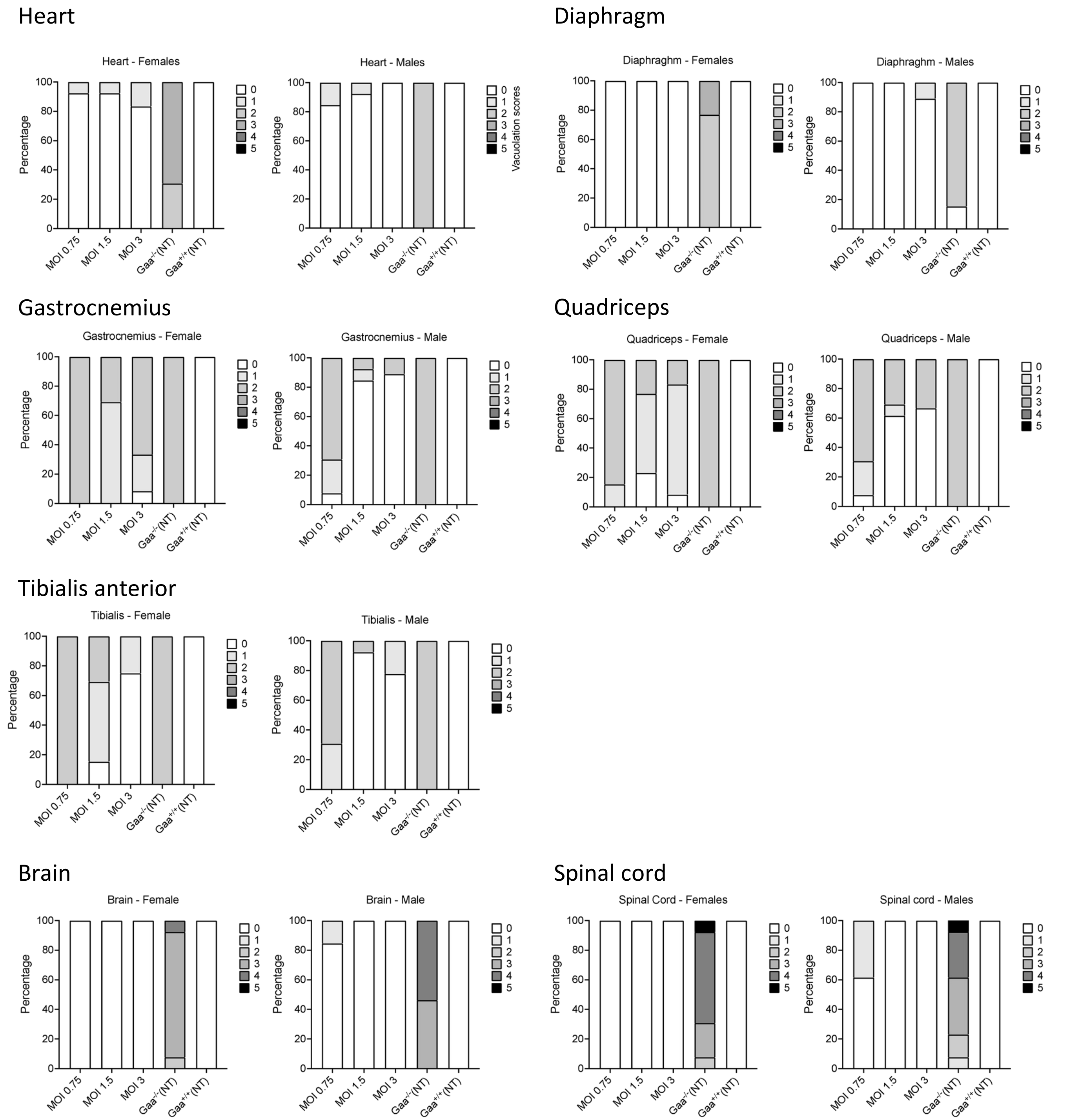


**Supplementary figure S4.** Rescue of pathology of tissues: Vacuolation scores in heart, and skeletal muscles diaphragm, gastrocnemius, quadriceps and tibialis anterior. CNS tissues included brain and spinal cord. Mouse tissue was analyzed at 8 months after transplantation (N = 9-13).

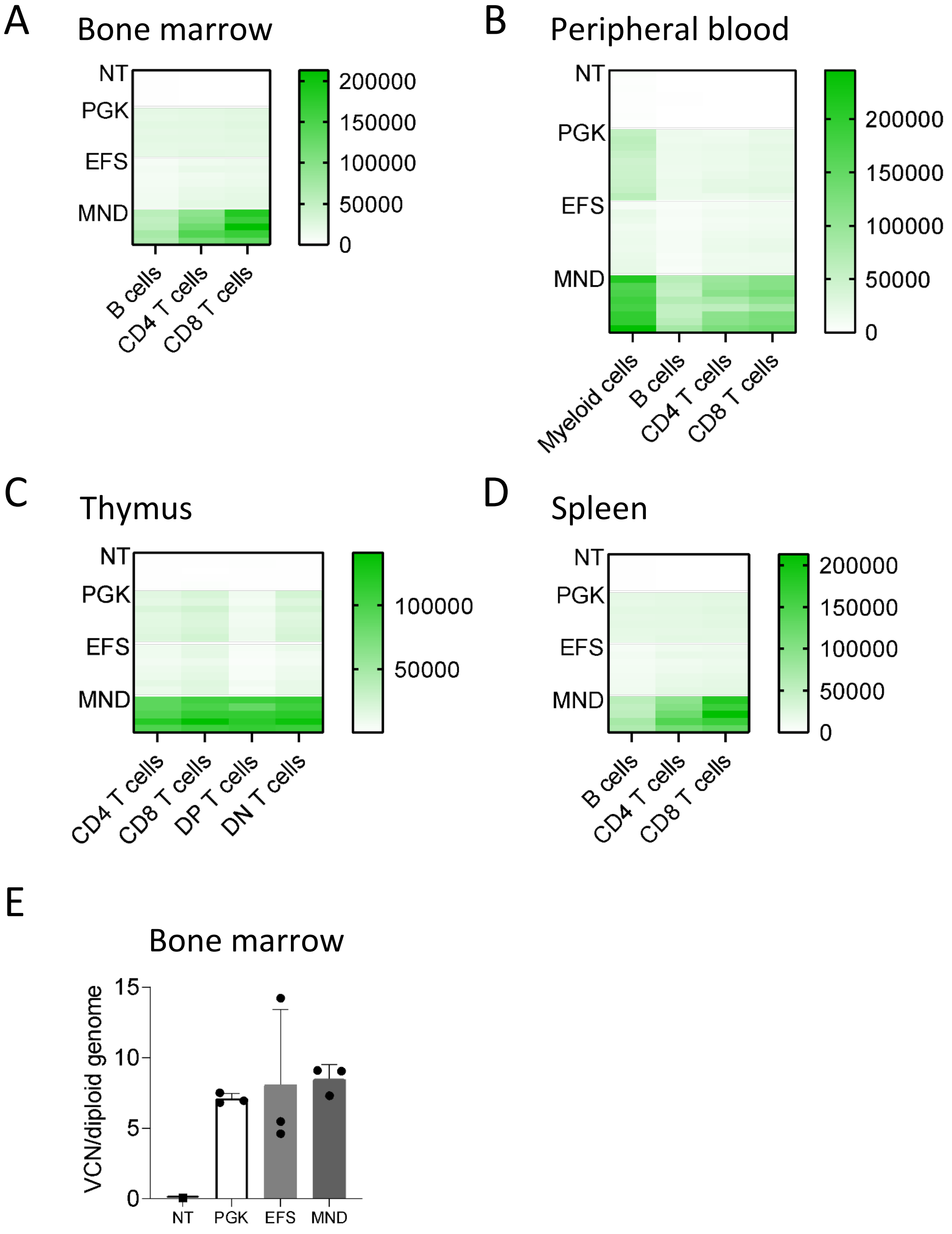


**Supplementary figure S5.** Promoter comparison in hematopoietic cells of GFP lentiviral vector treated mice. Mean fluorescence intensity (MFI) in hematopoietic tissues at 16 weeks after transplantation (n = 5 -8) in (A) bone marrow, (B) peripheral blood leukocytes, (C) thymus and (D) spleen. GFP MFI was assessed in different hematopoietic cell subsets, including myeloid, B cells and CD3+CD4+ and CD3+CD8+ T cells. In thymus double positive (DP) and double negative (DN) were assessed. (E) VCN in bone marrow at week 16 (n = 1 - 3). PGK = phosphoglycerate kinase promoter, EFS = elongation factor 1 alpha short promoter, VCN = vector copy number.

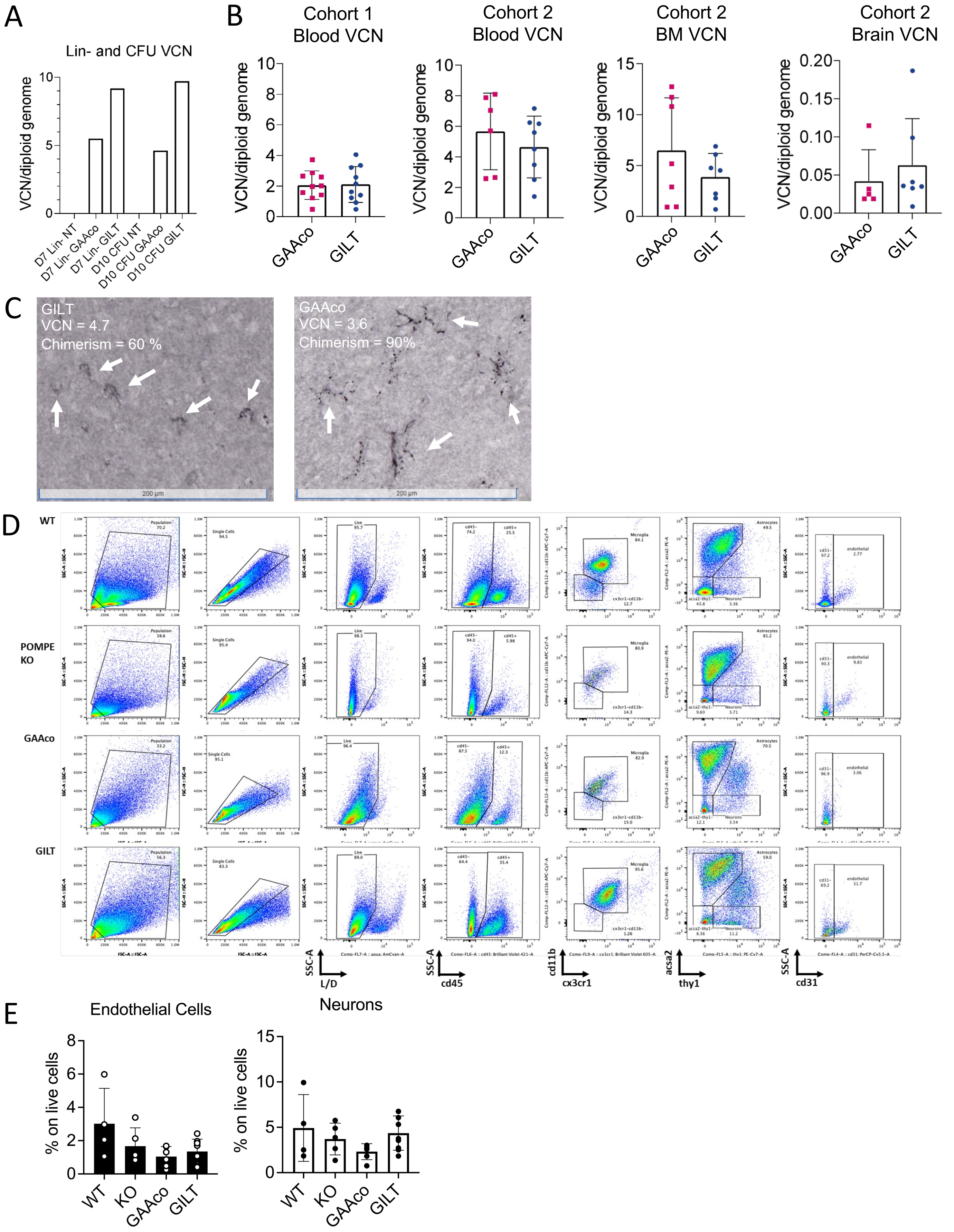


**Supplementary figure S6.** Transgene product and cell populations in brain of gene therapy treated *Gaa*$^{-/-}$ mice. (A) VCN of lentiviral vector SFFV promoter driven *GAAco* or *GILT* transgenes determined in bulk cultured transduced Lin- cells and combined CFU cultures. (B) VCN analysis in two cohorts showing similar VCN between GAAco and GILT groups (N= 5-8). (C) Immunohistochemistry staining for GAA protein in brain in cohort 1. Arrows indicate GAA positive cells. (D) FACS gating strategy for isolation of astrocytes, oligodendrocytes, neurons and oligodendrocytes from cohort 2. (E) Unaffected cell populations in the brain of *Gaa*$^{-/-}$ mice. Percentage endothelial cells and neurons (N=4-7). Samples were collected at at 32-weeks after transplant. SFFV = spleen focus forming virus promoter.

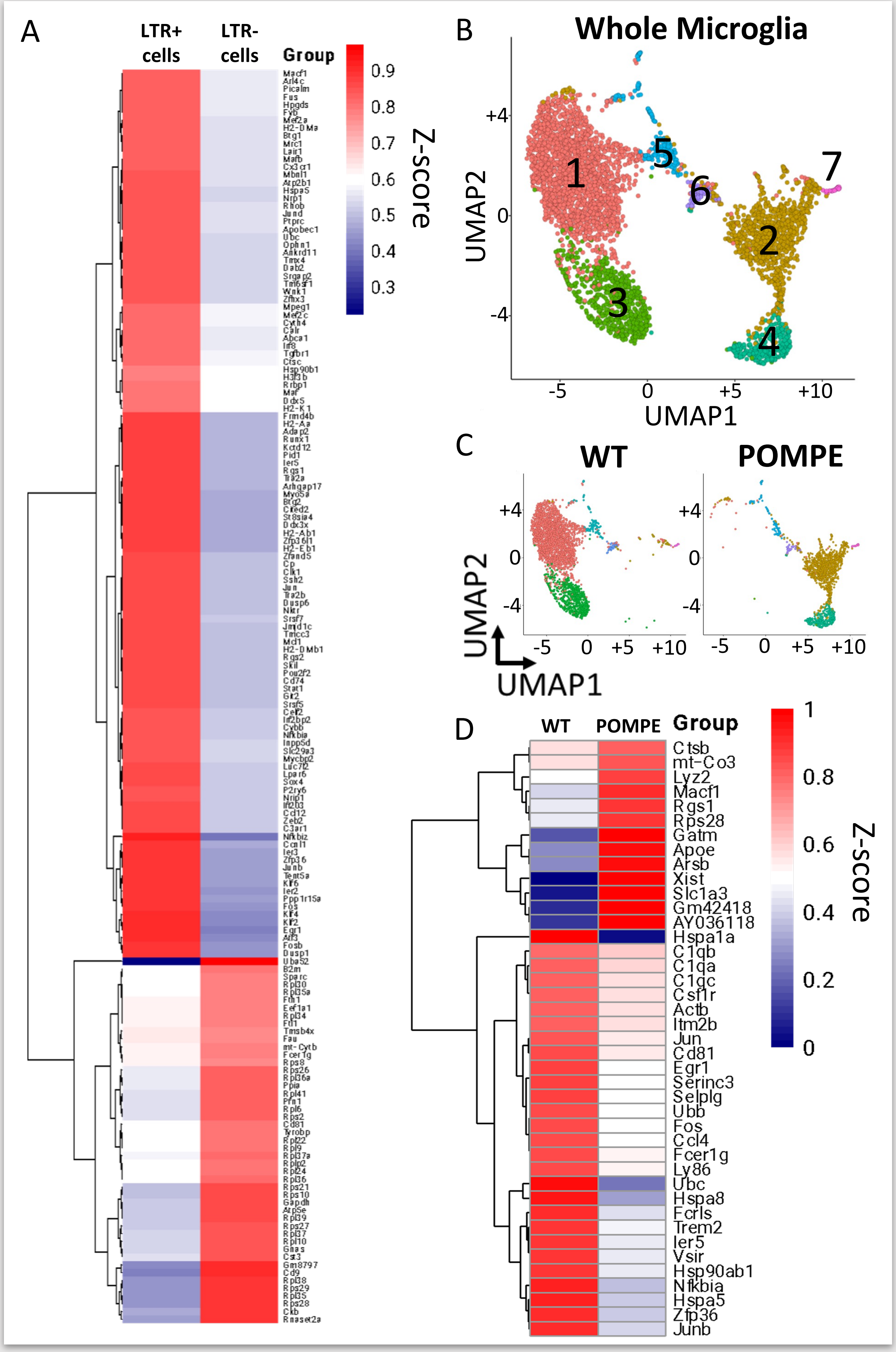
A
LTR+ cells
LTR- cells
Group
0.9
0.8
0.7
0.6
0.5
0.4
0.3
Z-score
B
Whole Microglia
UMAP2
UMAP1
+4
0
-4
-5
0
+5
+10
1
2
3
4
5
6
7
C
WT
POMPE
UMAP2
UMAP1
+4
0
-4
-5
0
+5
+10
D
WT
POMPE
Group
Ctsb
mt-Co3
Lyz2
Macf1
Rgs1
Rps28
Gatm
Apoe
Arsb
Xist
Slc1a3
Gm42418
AY036118
Hspa1a
C1qb
C1qa
C1qc
Csf1r
Actb
Itm2b
Jun
Cd81
Egr1
Serinc3
Selplg
Ubb
Fos
Ccl4
Fcer1g
Ly86
Ubc
Hspa8
Fcrls
Trem2
Ier5
Vsir
Hsp90ab1
Nfkbia
Hspa5
Zfp36
Junb
1
0.8
0.6
0.4
0.2
0
Z-score

**Supplementary figure S7.** Additional single-cell RNA sequencing analyses including data from microglial cells isolated from WT control and *Gaa*$^{-/-}$ mice. (A) Heatmap showing the average expression of genes specifically expressed in LTR+ vs LTR- cells from GILT and GAAco treated mice from the single-cell transcriptional profiling of Figure 5. (B) UMAP plot generated from a total of 4,135 single microglia cells isolated from WT control and *Gaa*$^{-/-}$ mice (n = 4 mice, 2 mice per group) clustered based on transcriptional similarities. The gene signatures relative to each cluster are listed in supplementary table S2. (C) UMAP plots showing single cell distribution in the WT (left plot) or *Gaa*$^{-/-}$ mice (right plot). (D) Heatmap showing average expression values of differentially expressed genes between WT control and *Gaa*$^{-/-}$ microglia cells.

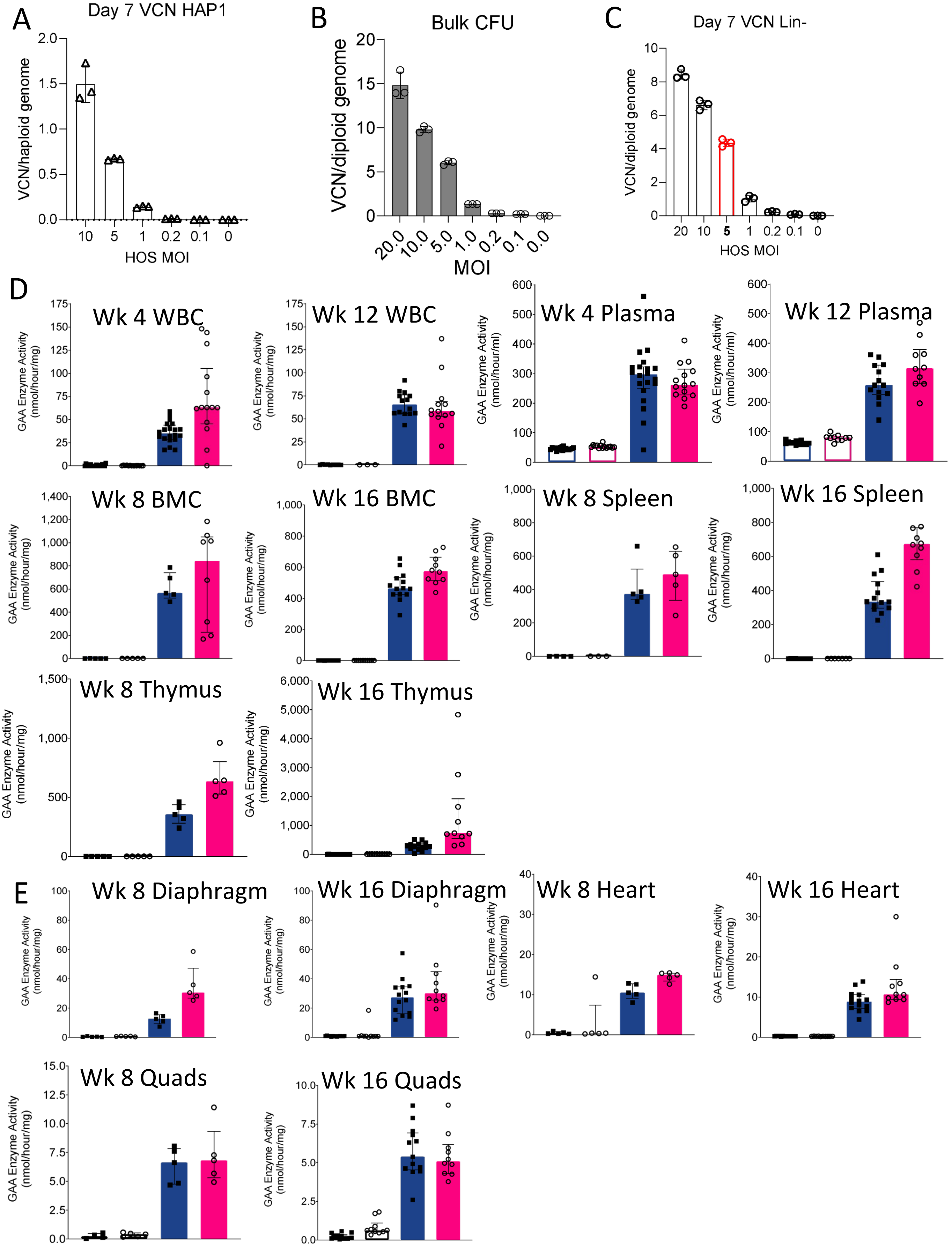


**Supplementary figure S8.** Successful engraftment of GILT transduced Lin- cells in Busulfex conditioned *Gaa*$^{-/-}$ mice. (A) Vector titration on HOS cells and subsequent VCN analysis (n = 3 per group) (B) Vector titration on Lin- cells and VCN analysis in bulk CFU suspensions (n = 3 per group). (C) Vector titration and VCN analysis by qPCR on Lin- cell suspensions (n = 3 per group). (D) GAA enzyme activity in peripheral blood and hematopoietic tissues (N = 5-14). (E) GAA enzyme activity in heart and skeletal muscles (N = 5-14). Blue = males and Pink = females.

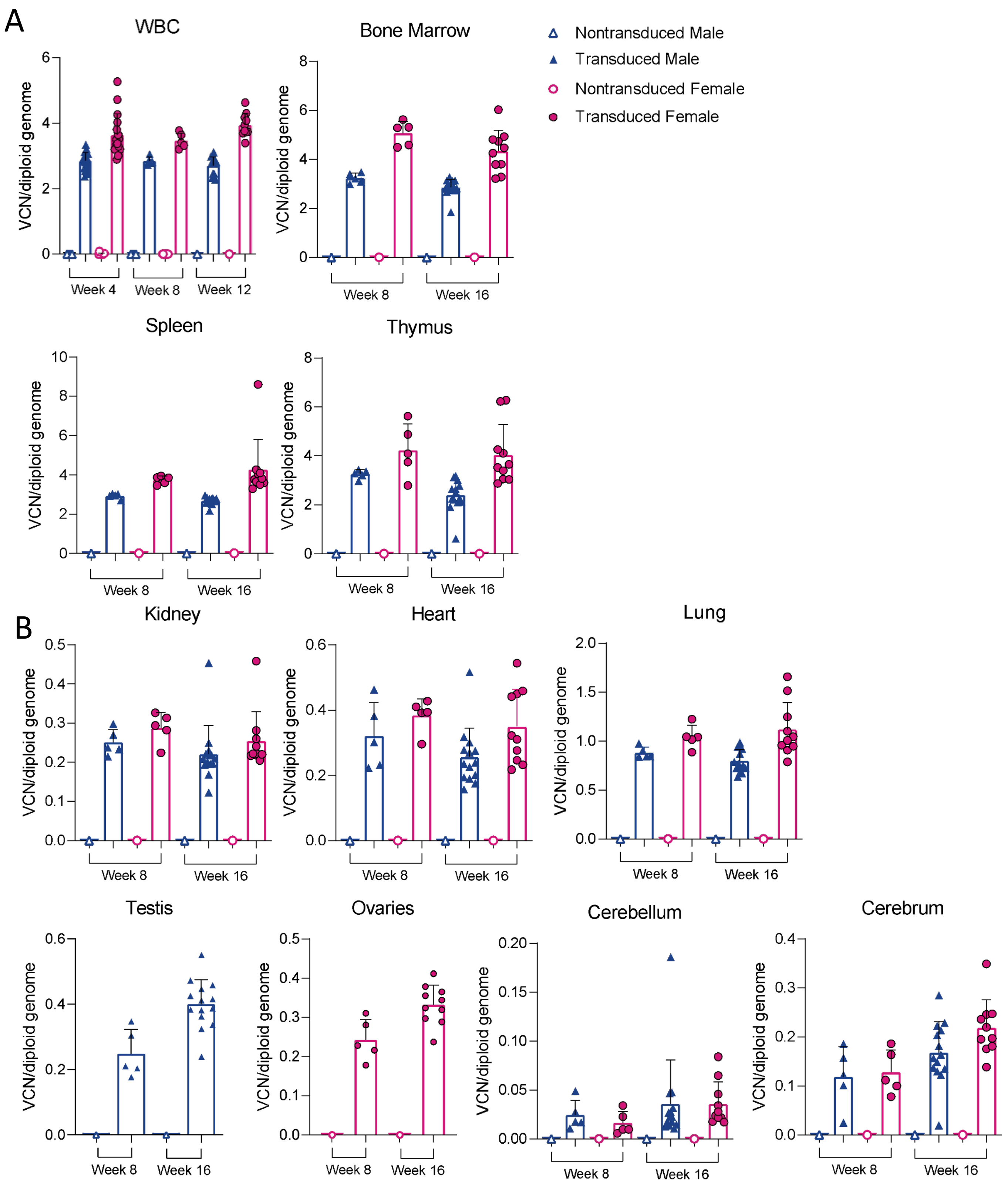


**Supplementary figure S9.** Vector copy numbers in Busulfex conditioned mice transplanted with genetically modified Lin- cells.
(A) VCN analysis in hematopoietic tissues (N = 1-14). (B) VCN in peripheral tissues including brain and reproductive organs (N = 1-14). Blue = males and Pink = females.

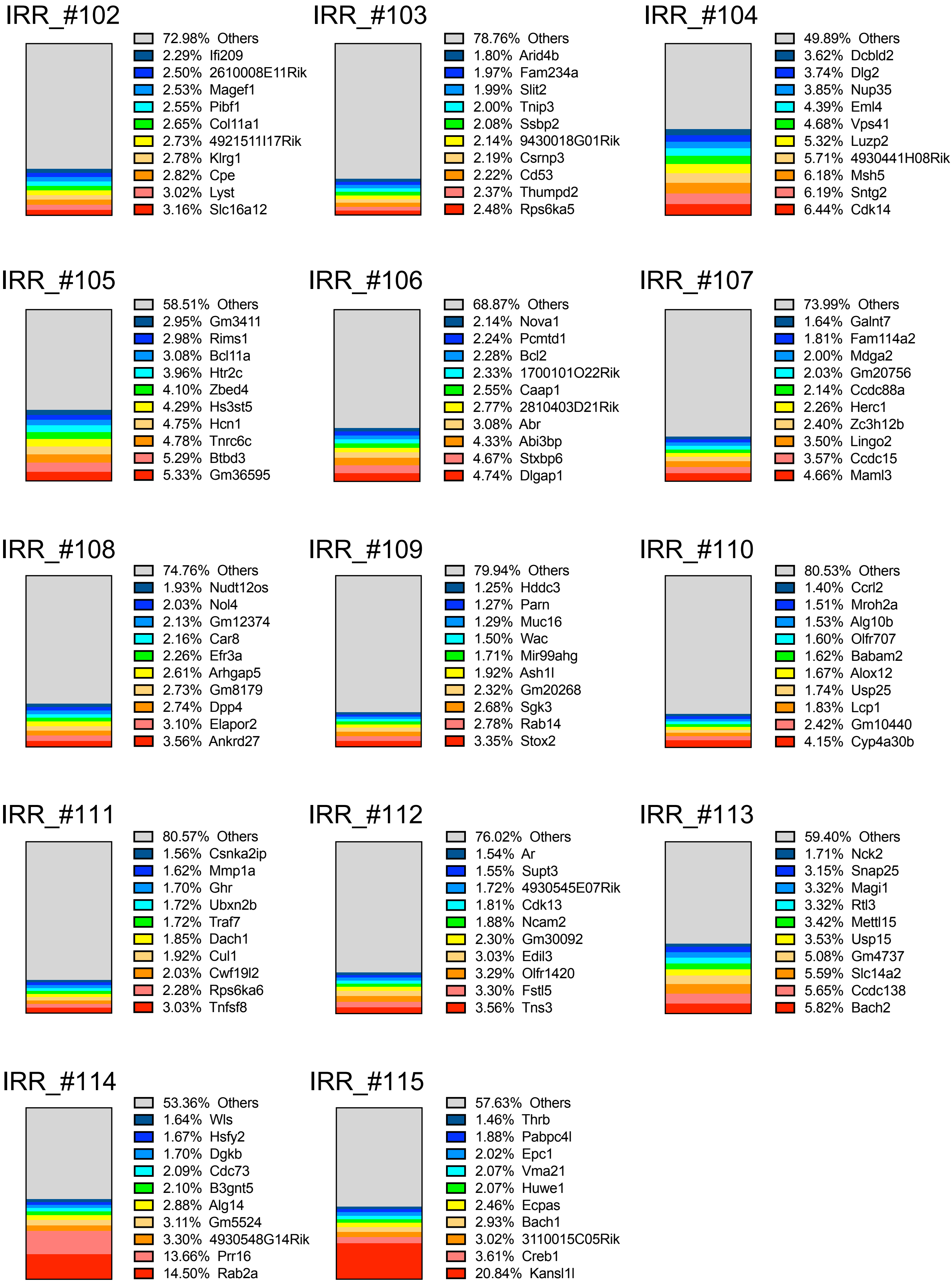


**Supplementary figure S10.** Top 10 integration sites of the of GILT and GFP treated mice.

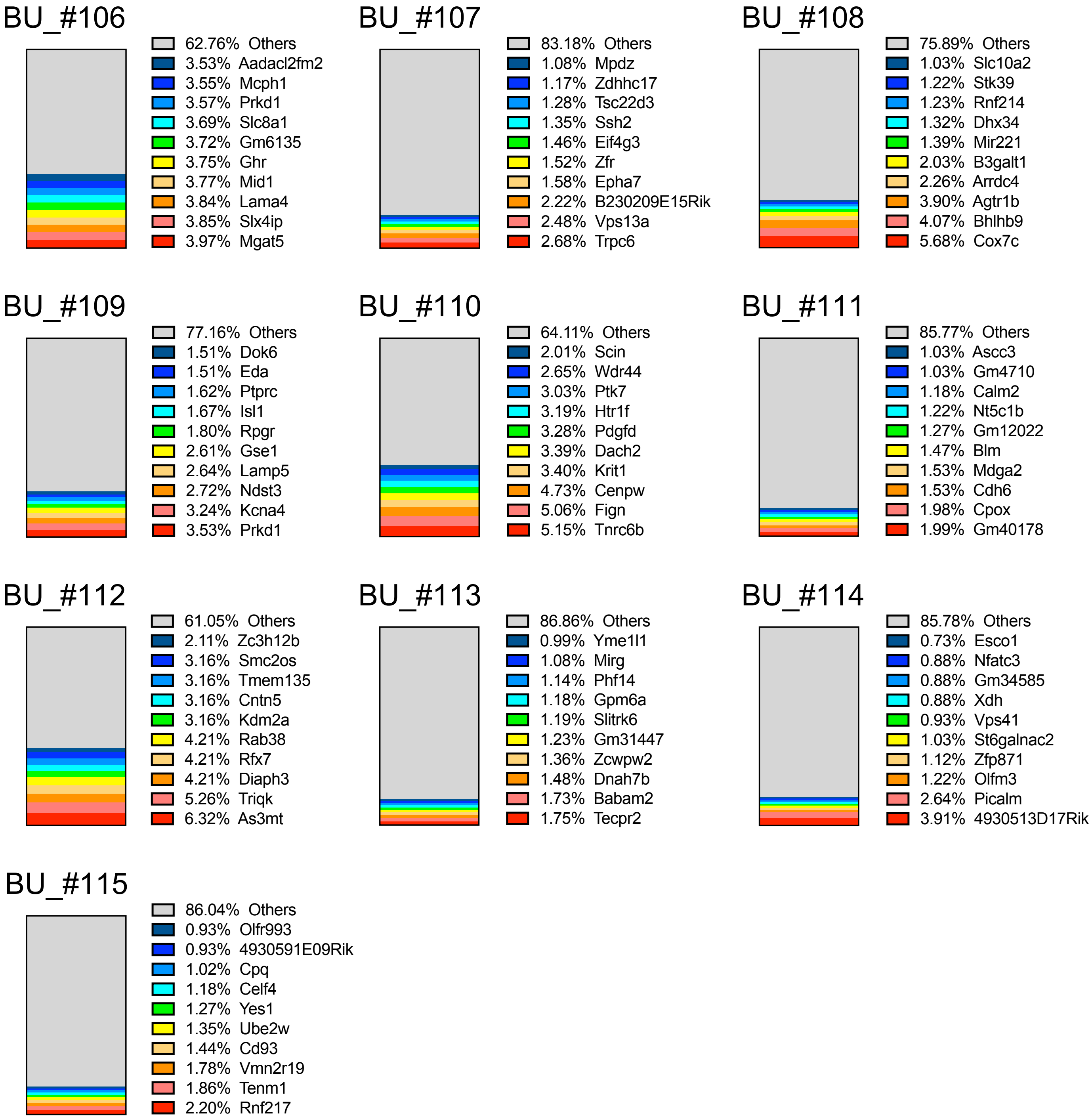


**Supplementary figure S10 continued.** Top 10 vector integration sites of the GILT and GFP treated mice.

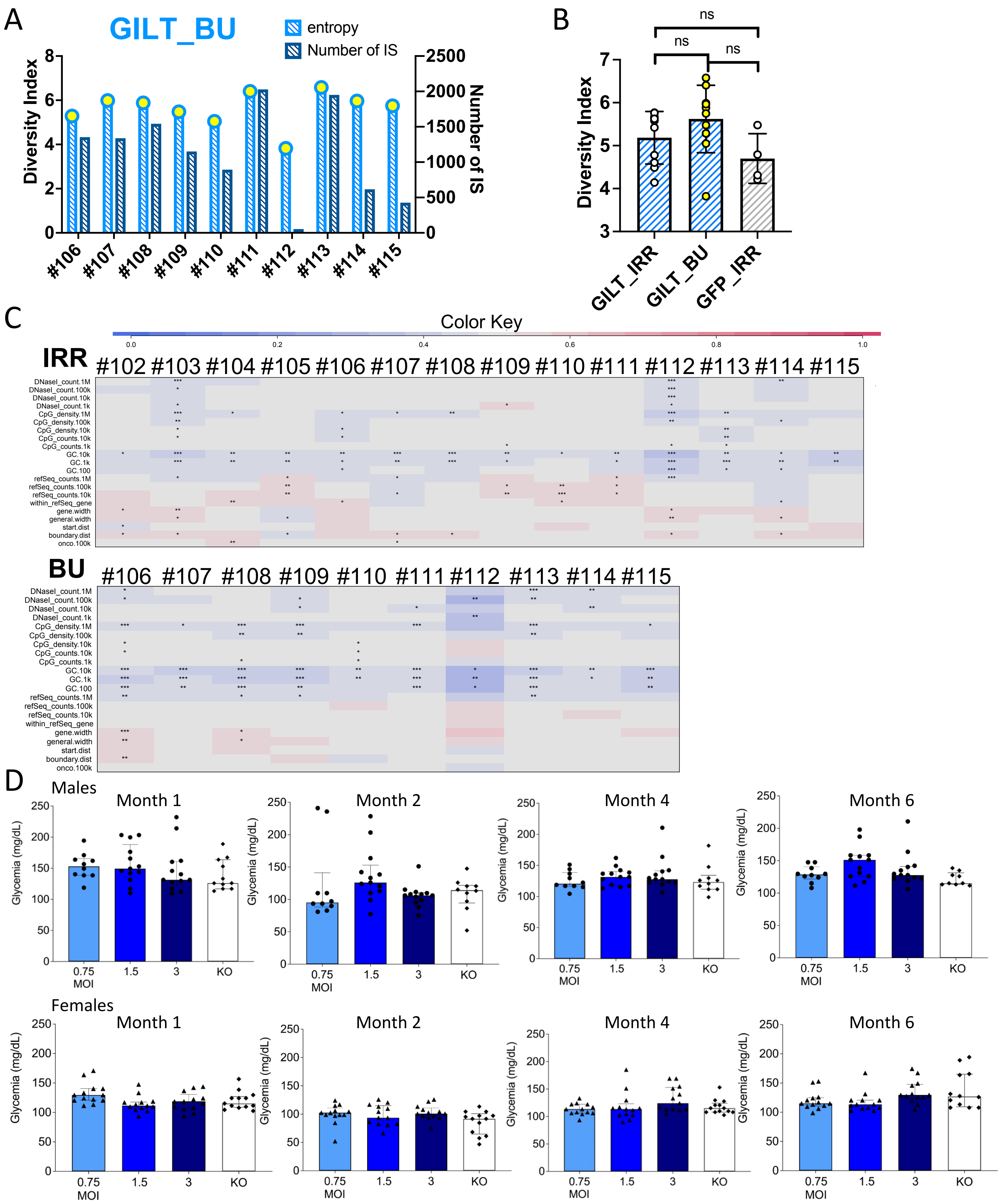


**Supplementary figure S11.** Safety assessment in GILT-vector treated mice.

(A) ) Busulfex (Bu) study GILT-group with Shannon diversity index and polyclonality at 16-weeks after transplantation. (B) Diversity index group comparison. (C) Genomic heatmap analysis of GILT-vector treated animals (102-115), or GILT-vector animals of the Bu study (106-115). The colored tiles (values of ROC areas) indicate the intensity and direction of any departures from the distribution of random controls for each feature in each integration site data set. A random distribution is represented by a ROC value of 0.5 and a grey color tiles as seen in the middle of the Color Key. A blue color indicates an under-representation, a red color an over-representation of integrations close to certain features. (D) Glucose levels over time in male and female mice. Glucose levels were measured at month 1, 2, 4 and 6. Mice were irradiation conditioned. Significant differences were not observed between groups ( N = 10-13).

**Supplementary table S1.** Abbreviation definition echocardiography

Short Axis of the heart in M-mode
**IVSd**, interventricular septum in diastole
**IVSs**, interventricular septum in systole
**LVIDd**, left ventricular internal diameter in diastole
**LVPWd**, left ventricular posterior wall in diastole
**LVIDs**, left ventricular internal diameter in systole
**LVPWs**, left ventricular posterior wall in systole
**LVmass**, left ventricular mass
**LVmass index**, left ventricular mass indexed by body weight
**LVVd**, left ventricular volume in diastole
**LVVs**, left ventricular volume in systole

**FS**, fractional shortening
**EF**, ejection fraction
**IVCT**, isovolumic contraction time

**Supplementary table S2.** Single cell RNA sequencing microglia gene signatures

A

| Cluster 1 | Cluster 2 | Cluster 3 | Cluster 4 | Cluster 5 |
|---|---|---|---|---|
| Fos | Fos | Fos | Rps29 | Rps29 |
| Jun | Jun | Jun | Rps28 | Rps28 |
| Fosb | Fosb | Fosb | Uba52 | Uba52 |
| Cx3cr1 | Cx3cr1 | Cx3cr1 | Rps21 | Rps21 |
| Mafb | Mafb | Mafb | Rpl38 | Rpl38 |
| Zeb2 | Zeb2 | Zeb2 | Rpl39 | Rpl39 |
| Tmem119 | Tmem119 | Tmem119 | Rpl37 | Rpl37 |
| Lair1 | Lair1 | Lair1 | Rpl10 | Rpl10 |
| Mrc1 | Cd74 | Qk | Rps27 | Rps27 |
| Ubc | H2-Eb1 | Son | Rps10 | Rps10 |
| Adap2 | H2-Aa | Pnisr | Rpl41 | Rpl41 |
| Nrp1 | H2-Ab1 | Srrm2 | Rpl37a | Rpl37a |
| Stab1 | H2-DMb1 | Atp6v0b | Tyrobp | Tyrobp |
| C3ar1 | Stat1 | Ogt | Selenop | Selenop |
| Rhoh | Ccl12 | Hnrnpa2b1 | Timp2 | Timp2 |
| Slc29a3 | H2-K1 | Chd9 | Fcer1g | Fcer1g |
| Olfml3 | Ciita | Tcf4 | B2m | B2m |
| Klf4 | H2-Q7 | Rbm25 | Cd63 | Cd63 |
| Csf1r | H2-D1 | Snrnp70 | Selenof | Selenof |
| Inpp5d | Cd52 | Nktr | Ubb | Ubb |
| Picalm | Irf1 | Arglu1 | Rpl28 | Rpl28 |
| Tra2b | Fxyd5 | Ttc14 | Ctss | Ctss |
| Ttc28 | Fgl2 | Rbm39 | Ctsd | Ctsd |
| Mertk | Ifitm3 | Mtdh | Lamp1 | Lamp1 |
| Slc15a3 | Tap1 | Zbtb20 | Hmgb1 | Hmgb1 |
| Mgat4a | H2-T23 | Sf3b1 | Serf2 | Serf2 |
| Irf2bp2 | Ctsc | Lpar6 | Hexb | Hexb |
| Ptgs1 | Mycbp2 | Arhgap5 | Cd9 | Cd9 |
| Tgfbr2 | Samhd1 | Gpr34 | Apoe | Sparcl1 |
| Cd33 | Apobec3 | Sat1 | | C1qb |
| Ccnd2 | Lilrb4a | | | Cd81 |
| Slc3a2 | Slfn2 | | | Ldhb |
| Dock4 | Psmb9 | | | C1qc |
| Pde3b | Ccr5 | | | Actb |
| P2ry6 | Ifi204 | | | Sparc |
| Ubash3b | Ly6e | | | Aldoa |
| Marcks | Ifi30 | | | Cdc42 |

B

| Cluster 1 | Cluster 2 | Cluster 3 | Cluster 4 | Cluster 5 | Cluster 6 | Cluster 7 |
|---|---|---|---|---|---|---|
| Hexb | Apoe | Mef2a | Mef2a | Cybb | Clec10a | Nrxn1 |
| Trem2 | Ctsb | Jun | Ptprc | Ccr2 | Mrc1 | Timp3 |
| P2ry12 | Cd63 | | | H2-Aa | Cd163 | Aldoc |
| Csf1r | Tyrobp | | | Clec12a | Ccl7 | Tmem47 |
| Cx3cr1 | | | | Cd74 | Ifitm2 | Apoe |
| Tmem119 | | | | S100a6 | Apoe | |
| | | | | Ifitm3 | Cybb | |
| | | | | Il1b | Ccl2 | |
| | | | | Ifitm2 | Ctsb | |
| | | | | Cxcl2 | Cd14 | |
| | | | | Tmem176a | Cd63 | |
| | | | | Tmem176b | Tmem176b | |
| | | | | Cd14 | Tyrobp | |
| | | | | Apoe | Cd68 | |
| | | | | Ptprc | | |
